# Macrophage-derived insulin/IGF antagonist ImpL2 regulates systemic metabolism for mounting an effective acute immune response in *Drosophila*

**DOI:** 10.1101/2020.09.24.311670

**Authors:** Gabriela Krejčová, Adam Bajgar, Pavla Nedbalová, Julie Kovářová, Nick Kamps-Hughes, Helena Zemanová, Lukáš Strych, Tomáš Doležal

**Affiliations:** University of South Bohemia, Czech Republic, United States; Biology Centre CAS, Czech Republic, United States; University of Oregon, United States

**Keywords:** macrophage, immuno-metabolism, Drosophila, immunity, bacterial infection, Streptococcus, ImpL2, IGFBP7, energy mobilization, Hif1α, aerobic glycolysis, macrophage polarization, insulin resistance, selfish immune system, adipose tissue remodeling, Foxo, infection-induced insulin resistance, Listeria, Insulin/IGF antagonist, wasting, cachexia

## Abstract

In response to invading pathogens, macrophages metabolically polarize towards Hif1α-induced aerobic glycolysis, requiring increased supply of nutrients. Here, we show that in order to obtain sufficient resources, Drosophila macrophages release the insulin/IGF antagonist ImpL2, whose expression is regulated by Hif1α. ImpL2 remotely induces the release of lipids and carbohydrates from adipose tissue by reducing insulin signaling, followed by increased nutrient accumulation in activated immune cells. ImpL2 thus translates the metabolic requirements of immune cells into a systemic metabolic switch. Although these ImpL2 effects are essential during the acute immune response to streptococcal infection, they become maladaptive upon chronic infection by an intracellular pathogen. The relevance of our model to mammalian immunometabolism is demonstrated by the increased expression of the ImpL2 homolog IGFBP7 in human macrophages exposed to Streptococcus.

## Introduction

Macrophages represent the front line of defence against invading pathogens. Although the effectiveness of the immune response correlates with the number of immune cells (Nicholson and Nicholson 2008), their maintenance requires energy and their excessive activation can lead to a myriad of pathologies and metabolic disorders (Shattuck-Heidorn et al. 2016; Zmora et al. 2017). Animals have therefore evolved a strategy that allows them to maintain sufficient numbers of quiescent immune cells that can be rapidly activated in response to the detection of pathogen- or danger-associated molecular patterns (Kelly and O’Neill 2015). Numerous populations of sentinel macrophages wait for activating stimuli without presenting a substantial energy burden. As a consequence, macrophages depend on rapid and sufficient supply from external sources, making the acute phase of the immune response challenging for the whole organism (Newsholme et al. 1986).

In response to the recognition of an invading pathogen, macrophages must rapidly alter their metabolism to generate enough energy and precursors to support their bactericidal function. Bactericidal (M1) macrophages therefore substantially increase the rate of glycolysis and the pentose phosphate pathway and rewire their mitochondrial metabolism in a Hif1α-dependent manner (Van den Bossche, O’Neill, and Menon 2017). Such a metabolic setup resembles the Warburg effect, which was originally described as a unique metabolic program for cancer cells (Warburg 1925). We have recently shown that the metabolic polarization of macrophages is an ancient and evolutionarily conserved process, as *Drosophila* macrophages also undergo a Hif1α-triggered metabolic switch that is essential for their bactericidal function (Krejčová et al. 2019).

An adverse aspect of M1 polarization is that these cells require more energy and become functionally dependent on external sources of glucose, glutamine, and lipids. Therefore, macrophages release signals in response to their metabolic activation that regulate systemic metabolism, thus securing nutrient supply at the expense of other organs (Straub 2014). Such privileged behavior, in which macrophages usurp nutrients from other processes, is crucial for an effective immune response (Bajgar and Dolezal 2018). One factor, mediating such behavior, is extracellular adenosine, which links the current metabolic state of activated immune cells to the systemic mobilization of carbohydrates that serve as a resource for immune defense (Bajgar et al. 2015; Bajgar and Dolezal 2018). However, this response is very complex and we assume the existence of other signaling factors with an analogous function.

To discover other signaling factors that are released by activated immune cells and regulate systemic metabolism, we sought inspiration from neoplastic tumor research. This idea is based on the notion that tumors and activated immune cells share common features of their cellular metabolism, as both utilize aerobic glycolysis triggered by Hif1α (Biswas and Mantovani 2012; Nagao et al. 2019). They also share an impact on systemic metabolism, as both cancer and sepsis patients exhibit a phenotype similar to the wasting caused by cytokine-induced insulin resistance (Dev, Bruera, and Dalal 2018). Although insulin resistance is mostly studied as a pathological condition, its evolutionary conservation indicates that it must carry an adaptive physiological function (Soeters and Soeters 2012; Odegaard and Chawla 2013). We therefore hypothesized that activated immune cells could release the same factors as tumor cells, but with a beneficial role for the acute response as opposed to cancer-induced cachexia.

In this study, we focus on the insulin/IGF antagonist ImpL2 (Imaginal morphogenesis protein-Late 2), which is released by neoplastic tumor cells, to suppress insulin signaling via binding to Drosophila insulin-like peptides, thereby causing energy wasting (Alee 2011; Arquier et al. 2006; Honegger et al. 2008; Kwon et al. 2015; Figueroa-Clarevega and Bilder 2015). In addition to its production by neoplastic tumors, ImpL2 is known to be released by lipid-overloaded macrophages (Morgantini et al. 2019), as well as by other cells employing Hif1α activity-dependent metabolic programs in *Drosophila* (Alee 2011; Owusu-Ansah, Song, and Perrimon 2013; Kwon et al. 2015; Figueroa-Clarevega and Bilder 2015). We therefore decided to test the role of ImpL2 as a macrophage-derived signaling factor that may be responsible for nutrient mobilization during the acute phase of immune response to bacterial infection.

Here we show that activated macrophages produce ImpL2 in a Hif1-dependent manner during the acute phase of infection, resulting in Foxo-mediated changes in adipose tissue metabolism. As an outcome of ImpL2 action, we observed increased titers of circulating carbohydrates and lipids and their accumulation in macrophages. ImpL2 release by macrophages is necessary for resistance to streptococcal infection. In contrast to this beneficial role in fighting extracellular pathogens, the effects of ImpL2 are maladaptive in response to intracellularly growing *Listeria*. Conservation of ImpL2 function between insects and mammals is indicated by increased expression of the ImpL2 homolog IGFBP7 in human macrophages exposed to streptococci.

## Results

### ImpL2 expression increases in immune-activated macrophages in a Hif1α-dependent manner

To test the potential role of ImpL2 during infection, we first monitored its expression profile during the acute phase of *Streptococcus pneumoniae* infection. ImpL2 expression increased significantly in infected flies compared to PBS-injected controls, as early as 3 hours post-infection (hpi), reaching up to a threefold increase in expression 21 hpi on an organismal level (Figure 1A). To identify the tissues responsible for the infection-induced rise in ImpL2 expression, we employed the Gal4 driver specific for the *ImpL2-RA* transcriptional variant (Bader et al. 2013) to drive expression of the fluorescent marker UAS-mCherry. The pattern of ImpL2-positive cells resembled the characteristic distribution of hemocytes in adult flies (Figure 1B), and their number increased substantially upon infection (Figure 1B and C). The ImpL2-RA>mCherry marker clearly colocalized with hemocyte-specific antibody against the scavenger receptor Nimrod C1 (NimC1; Figure 1D). To verify that cells expressing ImpL2-RA>mCherry are macrophages, we injected flies with the phagocytic marker *S. aureus*-pHrodo-Green, and it indeed colocalized with these cells (Figure 1-figure supplement 1). Cells expressing ImpL2-RA>mCherry also actively recognized and engulfed *S. pneumoniae ex vivo* (Figure 1E). The evidence that ImpL2 is produced by macrophages during infection is further supported by the more than sixfold increase in expression in CrqGal4>UAS-GFP labeled (Clark et al. 2011) FACS sorted macrophages (Figure 1F).

**Figure 1.**
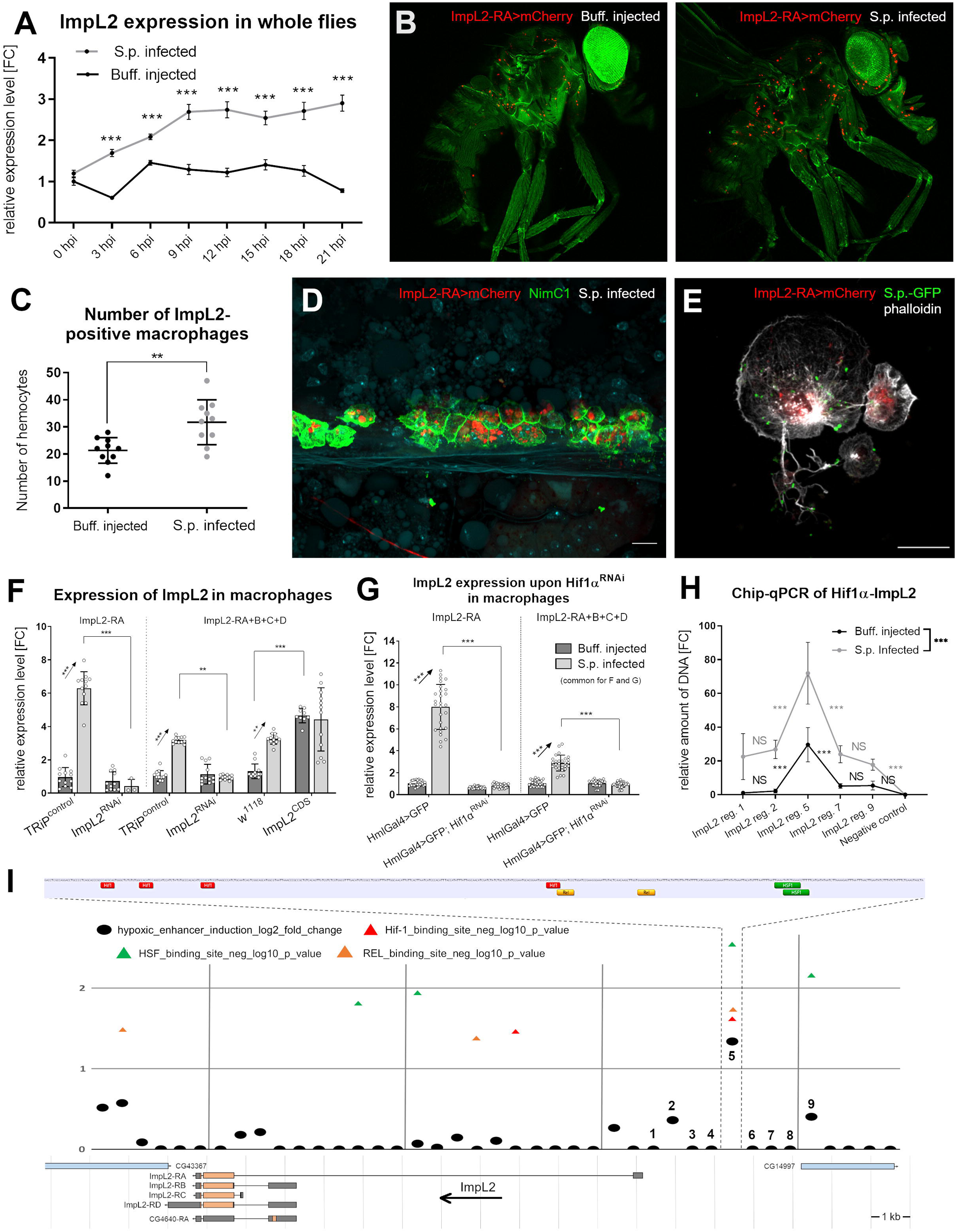
Streptococcal infection increases ImpL2 expression in a Hif1α-dependent manner in activated macrophages. (**A**) Expression of ImpL2 mRNA at whole-body level in flies (CrqGal4>GFP) infected with *S. pneumoniae* (S.p. infected) and control flies (Buff. injected) at various time points; results ompared by unpaired t test with Holm-Sidak method for multiple comparisons. (**B**) Representative confocal microscopy images of control (left) and infected (right) ImpL2-RA>mCherry individuals imaged at 24 hpi, from Z stack of 7 layers, autofluorescence in the green channel was used to visualize the fly’s body (see Figure 1-figure supplement 5 for images in color blind friendly pallete). (**C**) Number of ImpL2-positive macrophages in the thorax of control and infected adult flies (ImpL2-RA>mCherry) at 24 hpi; results compared by unpaired t test. (**D**) Confocal microscopy image of infected and dissected ImpL2-RA>mCherry bearing fly stained with anti-NimC1 antibody (green) depicting the expression of ImpL2 in hemocytes, from Z stack of 11 layers. The scale bar represents 10 µm. (**E**) Confocal microscopy image of ImpL2-RA>mCherry hemocyte actively interacting with GFP-labeled *S. pneumoniae* (S.p.-GFP) ex vivo. The scale bar represents 13 µm. (**F**) Expression of ImpL2 mRNA (using primers targeting all transcript variants) in hemocytes isolated from Buff. injected and S.p. infected flies with macrophage-specific ImpL2 knockdown (ImpL2^RNAi^), overexpression (ImpL2^CDS^) and their respective controls (TRiP^control^, w1118) at 24 hpi (right) and gene expression of ImpL2-RA transcript in hemocytes isolated from Buff. injected and S.p. infected flies with hemocyte-specific ImpL2 knockdown (ImpL2^RNAi^) and its respective control (TRiP^control^) at 24 hpi (left), documenting the efficiency of ^RNAi^ fly line used in this manuscript. Results compared by 2way ANOVA Tukey’s multiple comparisons test. (**G**) Gene expression of ImpL2-RA (left) and all ImpL2 transcript variants (right) in hemocytes isolated from Buff. injected and S.p. infected control (HmlGal4>GFP) flies and flies with hemocyte-specific Hif1α knockdown at 24 hpi. Results compared by 2way ANOVA Tukey’s multiple comparisons test. (**H**) Relative amount of selected ImpL2 genomic regions (visualized in I) bound by the transcription factor Hif1α in Buff. injected ans S.p. infected flies. Selected genomic region of S-adenosylmethionin synthetase was used as a negative control. Results compared by 2way ANOVA Tukey’s multiple comparisons test. (**I**) In silico analysis of hypoxic enhancer activity by 500 bp bins at the ImpL2 locus and visualization of ImpL2 transcript variants (Flybase.org). Each black dot plots a log2 fold change (y-axis) of the difference in the randomer tag counts mapped to the 500-bp bin between normoxic and hypoxic conditions. The triangles show the position and negative log10 p-value (y-axis) of multiple hypoxic (Hif1α), immune (Rel) and stress (HSF) response transcription factor binding sites. The close up shows distribution and clustering of individual response elements for the most significant bin. In A, F and G, expression levels normalized against rp49 are reported as fold change relative to levels of ImpL2 (using primers targeting all transcript variants) and ImpL2-RA respectively, in Buff. injected controls, which were arbitrarily set to 1. The individual dots represent biological replicates with line/bar showing mean ± SD, asterisks mark statistically significant differences (*p<0.05; **p<0.01; ***p<0.001). Hif1α, hypoxia-inducible factor 1 α; Rel, relish; HSF, heat shock factor.

As there are three alternative transcriptional start sites for ImpL2 (Figure 1I), we analyzed the expression pattern of each isoform at 24 hpi. Among all analyzed isoforms, the expression of *ImpL2-RA* was the highest of all transcriptional variants in macrophages and was hardly detectable in fat body or muscles (Figure 1-figure supplement 2). While the *ImpL2-RB+D* forms were the most abundant variants in adipose tissue, their expression was five times weaker than *ImpL2-RA* expression in macrophages (see different y-axis scales in Figure 1-figure supplement 2). In addition, expression of the *ImpL2-RA* isoform increased more than sixfold in macrophages, the most significant increase in expression of the transcripts in response to infection (Figure 1F and Figure 1-figure supplement 2). Thus, the *ImpL2-RA* transcriptional variant expressed in macrophages contributes significantly to the overall increase in ImpL2 expression observed at the onset of infection.

The metabolic switch in activated macrophages is regulated by the transcription factor Hif1α (Krejčová et al. 2019), which is known to be a potent regulator of ImpL2 expression in cells, which rely on anaerobic metabolism (Allee 2011; Li et al. 2013; Owusu-Ansah, Song, and Perrimon 2013). We therefore hypothesized that this same regulation could underlie the infection-induced increase in ImpL2 in activated macrophages. Macrophage-specific knockdown of Hif1α (for efficiency, see Figure 1-figure supplement 3) resulted in the inability of these cells to trigger the characteristic infection-induced expression of ImpL2, and this effect was particularly evident for the *ImpL2-RA* isoform (Figure 1G). We identified a cluster of four Hif1α binding sites upstream of the transcription start site of *ImpL2-RA* (Figure 1I). Moreover, this enhancer region shows the strongest hypoxic induction of all sequence surrounding the *ImpL2* gene (Kamps-Hughes et al. 2015 and Figure 1I), suggesting that Hif1α may directly drive *ImpL2-RA* transcription in macrophages. In-depth analysis of this 500bp region revealed the presence of eight immune and stress response elements clustered together (four hypoxia response elements, two Relish bindings sites, and two heat shock factor binding sites; Figure 1I and Figure 1-figure supplement 4). Chip-qPCR analysis revealed direct binding of Hif1α to this part of the *ImpL2-RA* promoter, and this interaction was further enhanced after infection (in Figure 1H denoted as region 5).

These experiments identified activated macrophages as prominent producers of ImpL2 in infected adult flies. The increase of ImpL2 production was associated with a Hif1α-induced metabolic switch in macrophages, through direct binding of Hif1α to the regulatory sequence of the *ImpL2-RA* isoform.

### Macrophage-derived ImpL2 is required for Foxo-mediated mobilization of fat body reserves

Activated macrophages release signaling factors that mobilize reserves to provide sufficient nutrients for the activated immune system (Bajgar and Dolezal 2018; Dolezal et al. 2019). Here we show that macrophages increase ImpL2 production during infection. As ImpL2 has previously been associated with reserve mobilization, leading to wasting in flies with experimentally induced neoplastic growth, we further tested the impact of macrophage-derived ImpL2 on systemic metabolism. To do so we employed *Drosophila* genetic tools to manipulate ImpL2 expression specifically in macrophages (Crq>Gal4) or specifically in macrophages temporally restricted to the adult stage (Hml>Gal4, Gal80^TS^). Using conventional ImpL2 knockdown (ImpL2^RNAi^) and overexpression (ImpL2^CDS^) constructs, we achieved significant changes in ImpL2 expression (Figure 1F). These experimental manipulations allowed us to either prevent infection-induced upregulation of ImpL2 in macrophages or to simulate the increase in ImpL2 expression in uninfected individuals (Figure 1F).

Both infection and overexpression of ImpL2 in macrophages significantly reduced triglyceride content in whole flies (Figure 2A), which was accompanied by dramatic changes in adipose tissue morphology (Figure 2B-E). The number of lipid droplets increased as their average size decreased (Figure 2B, C, and E), making the lipids more accessible for lipases. The overall area of adipose tissue occupied by lipid droplets was markedly smaller (Figure 2D). It should be emphasized that these infection-induced effects were suppressed by macrophage-specific ImpL2 knockdown (Figure 2A-E). Detailed lipidomic analysis of adipose tissue by mass spectrometry revealed that both infection and macrophage-specific overexpression of ImpL2 caused a proportional shift in lipid content from storage lipids (triglycerides) to polar lipid species (phosphatidylethanolamine, phosphatidylinositol), which participate in lipid mobilization and transport (Figure 2F). These effects were reduced by ImpL2 knockdown (Figure 2F). Similar effects were also observed for glycogen stores as both infection and overexpression of ImpL2 significantly reduced glycogen content in whole flies, while knockdown of ImpL2 abolished this effect (Figure 2G). This data thus indicate that macrophage-derived ImpL2 may serve as a mediator of cross-talk between the immune system and lipid metabolism in the adipose tissue.

**Figure 2.**
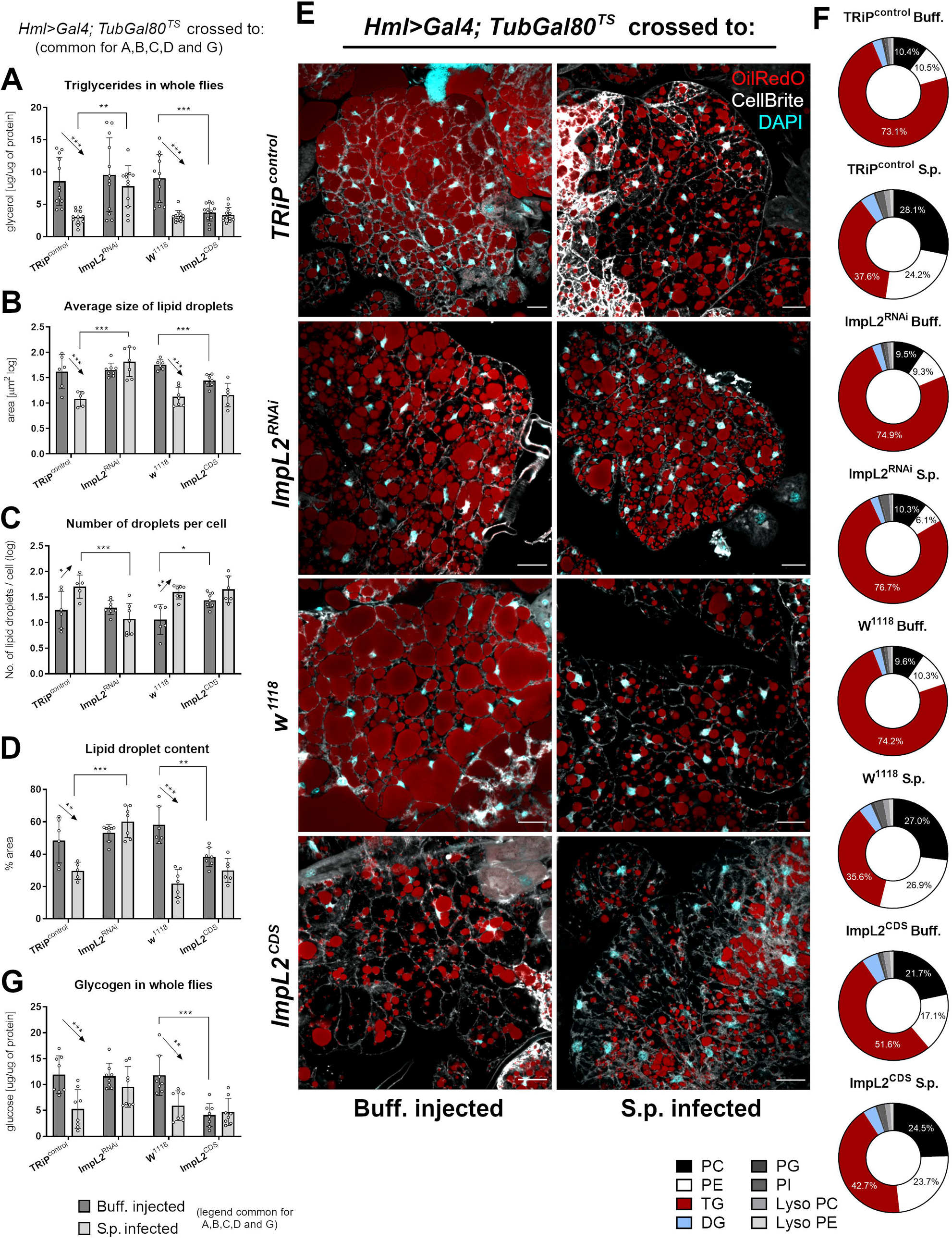
Macrophage-derived ImpL2 regulates changes in adipose tissue metabolism induced by infection. (**A**) Triglyceride concentration in Buff. injected and S.p. infected flies with macrophage-specific ImpL2 knockdown (ImpL2^RNAi^), overexpression (ImpL2^CDS^) and their respective controls (TRiP^control^, w1118) at whole-body level at 24 hpi. (**B-D**) Average size (**B**) and number (**C**) of lipid droplets (values log10-transformed) and percentage of area occupied (**D**) in the fat body of Buff. injected and S.p. infected flies with macrophage-specific ImpL2 knockdown (ImpL2^RNAi^), overexpression (ImpL2^CDS^) and their respective controls (TRiP^control^, w1118) at 24 hpi. (**E**) Representative confocal microscopy images of dissected fat body of Buff. injected and S.p. infected flies with macrophage-specific ImpL2 knockdown (ImpL2^RNAi^), overexpression (ImpL2^CDS^) and their respective controls (TRiP^control^, w1118) at 24 hpi, stained with OilRedO (red), DAPI (cyan) and CellBrite (white). The scale bar represents 20 µm. (**F**) Relative proportions of different lipid species in the fat body of Buff. injected and S.p. infected flies with macrophage-specific ImpL2 knockdown (ImpL2^RNAi^), overexpression (ImpL2^CDS^) and their respective controls (TRiP^control^, w1118) at 24 hpi. (**G**) Glycogen concentration in Buff. injected and S.p. infected flies with-specific ImpL2 knockdown (ImpL2^RNAi^), overexpression (ImpL2^CDS^) and their respective controls (TRiP^control^, w1118) at whole-body level at 24 hpi. Metabolite concentrations were normalized to the amount of proteins in each sample. Values are mean ± SD, asterisks mark statistically significant differences (*p<0.05; **p<0.01; ***p<0.001). Results in (**A-D** and **G**) were compared by 2way ANOVA Tukey’s multiple comparisons test. (**B-D**) were quantified from 8 confocal microscopy images for each genotype and treatment. Lipidomic analysis in (**F**) was performed in three biological replicates for each genotype and treatment. PC, phosphatidylcholine; PE, phosphatidylethanolamine; TG, triglycerides; DG, diglycerides; PG, phosphatidylglycerol; PI, phosphatidylinositol; LysoPC, Lyso-phosphatidylcholine; LysoPE, Lyso-phosphatidylethanolamine.

In addition to a substantially increased lipolytic and glycogenolytic programs, we also observed enhanced autophagy in the adipose tissue of infected flies bearing the Atg8a-mCherry reporter (Figure 3A), further implying an increase in catabolic metabolism. These infection-induced metabolic changes are consistent with the expression of many metabolic genes in adipose tissue. The expression of *MTP*, *apoLPP*, *apoLTP,* and *Bmm* genes associated with lipid mobilization, as well as *Atg1* and *Atg6* genes associated with autophagy, were upregulated in adipose tissue during infection (Figure 3B and Figure 3-figure supplement 1). Furthermore, their expression is under the control of macrophage-derived ImpL2. Indeed, knockdown of ImpL2 suppressed this effect, whereas overexpression of ImpL2 mimicked the response induced by infection (Figure 3B and Figure 3-figure supplement 1).

**Figure 3.**
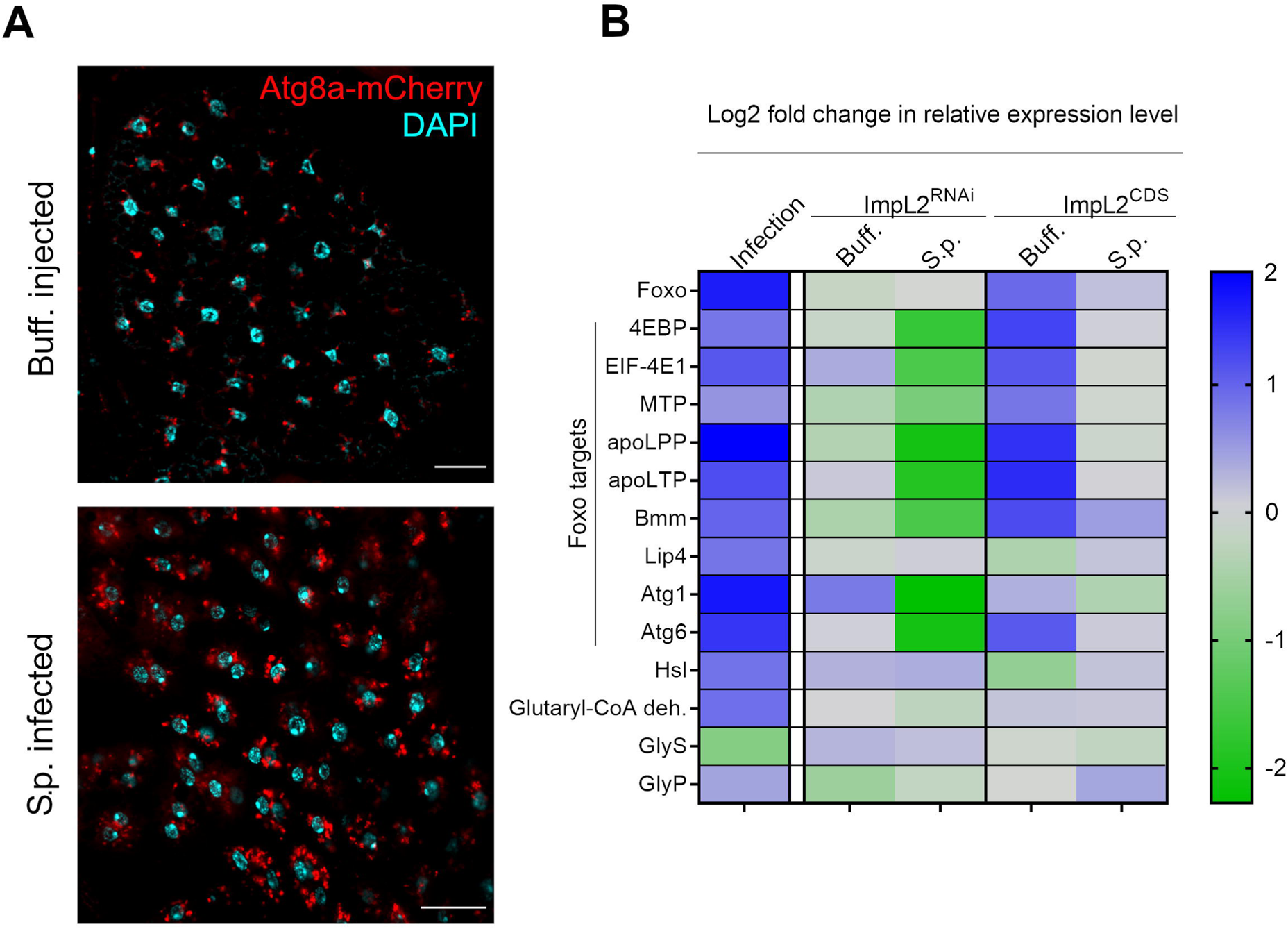
Infection-induced autophagy and changes in gene expression in adipose tissue. (**A**) Representative confocal microscopy images of fat body of control and infected flies with Atg8a-mCherry autophagy reporter (Atg8a, red; DAPI, cyan). The scale bar represents 20 µm. (**B**) Graphical representation of log2-fold change in mRNA expression of genes involved in insulin signaling, lipoprotein and lipid metabolism, glycogenolysis and autophagy in the fat body of Buff.-injected and S.p.-infected flies with macrophage-specific ImpL2 knockdown (ImpL2^RNAi^), overexpression (ImpL2^CDS^) and their respective controls (TRiP^control^, w1118) at 24 hpi. The first column (Infection) shows the log2-fold change in mRNA expression in S.p-infected flies compared to Buff.-injected control flies (average change for TRiP^control^ and w1118). The second to fourth columns represent the log2-fold change in mRNA expression compared to the corresponding control for knockdown (ImpL^RNAi^ to TRiP^control^) and overexpression (ImpL2^CDS^ to w1118) in either uninfected (Buff.) or S.p. infected (S.p.) flies. Foxo, forkhead box O; 4EBP, Thor; EIF-4E1, eukaryotic translation initiation factor 4E1; MTP, microsomal triacylglycerol transfer protein; apoLPP, apolipophorin; apoLTP, apolipoprotein lipid transfer particle; Bmm, brummer; Lip4, lipase 4; Hsl, hormone sensitive lipase; Glutaryl-CoA deh., Glutaryl-CoA dehydrogenase; GlyS, glycogen synthase; GlyP, glycogen phosphorylase; Atg1, autophagy-related gene 1; Atg6, autophagy-related gene 6.

Many of the aforementioned metabolic genes whose expression is altered by ImpL2 during infection (Figure 3B) are known Foxo targets, therefore we analyzed the impact of macrophage-derived ImpL2 on the subcellular localization of Foxo in adipose tissue. We found that Foxo displays nuclear localization upon infection, whereas it remained predominantly in the cytoplasm of adipocytes in uninfected flies (Figure 4). Macrophage-specific overexpression of ImpL2 recapitulated the infection-induced effect on Foxo nuclear localization, and conversely, knockdown of ImpL2 reversed this effect (Figure 4). These effects of ImpL2 on Foxo localization are in agreement with the expression of Foxo target genes (Figure 3B and Figure 3-figure supplement 1). To verify that the observed effects of macrophage-derived ImpL2 on the fat body are indeed mediated by Foxo, we tried to rescue the effect of ImpL2 overexpression in hemocytes by hypomorphic mutation of *Foxo*. The lipid droplet phenotype induced by ImpL2 overexpression was completely reversed by the heterozygous *foxo*^BG01018^ mutation (Figure 5A), demonstrating that the effects of macrophage-derived ImpL2 on fat body metabolism are indeed mediated by Foxo.

**Figure 4.**
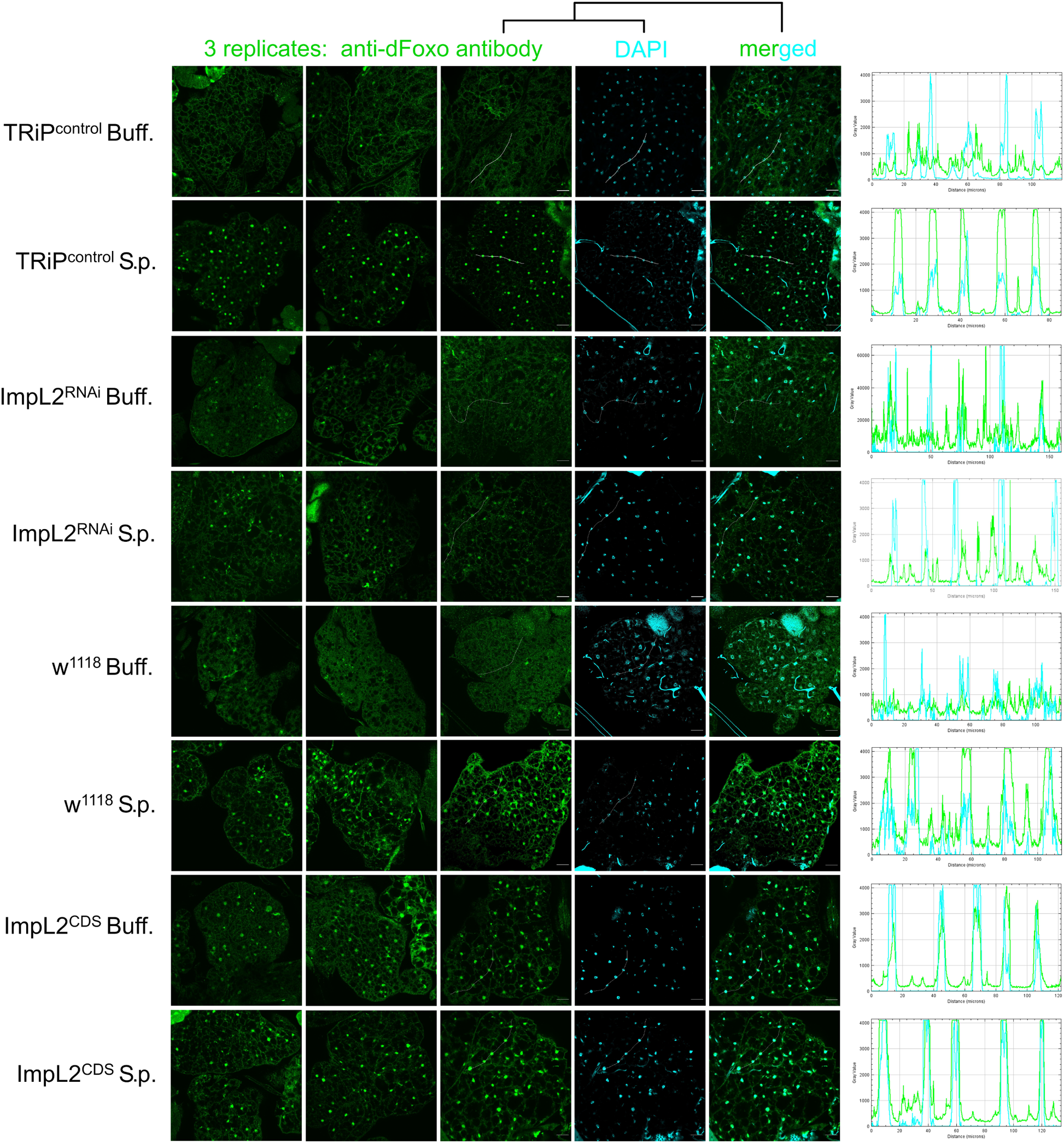
Macrophage-derived ImpL2 affects Foxo activity in adipose tissue. Representative confocal microscopy images of Foxo immunolocalization in the fat body of Buff. injected and S.p. infected flies with macrophage-specific ImpL2 knockdown (ImpL2^RNAi^), overexpression (ImpL2^CDS^) and their respective controls (TRiP^control^, w1118) at 24 hpi; anti-dFoxo antibody, green; DAPI, cyan. Histograms of Foxo cellular localization represent Foxo and DAPI signal intensity in sections indicated by a white freehand line. The scale bar represents 20 µm.

**Figure 5.**
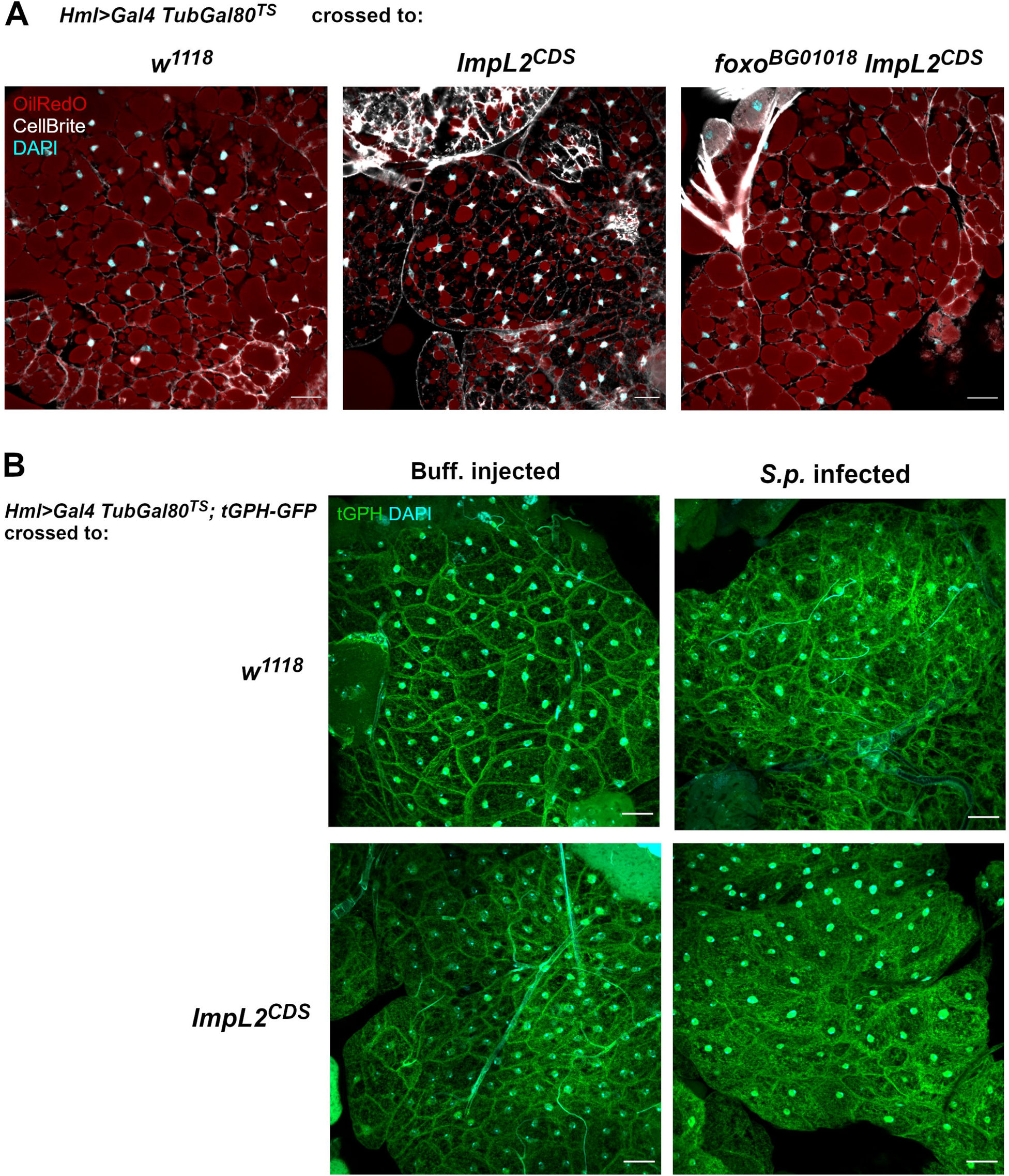
Macrophage-derived ImpL2 affects adipose tissue insulin sensitivity. (**A**) Representative confocal images of dissected adipose tissue of control flies (w1118), flies with macrophage-specific ImpL2 overexpression (ImpL2^CDS^), and flies with macrophage-specific ImpL2 overexpression (ImpL2^CDS^) and ubiquitous expression of hypomorphic variant of Foxo (foxoBG01018) stained by OilRedO (red), CellBrite (white) and DAPI (cyan) at 24 hpi. The scale bar represents 20 µm. (**B**) Representative confocal images of localization of the PI3K reporter tGPH in adipose tissue of Buff. injected and S.p. infected control flies (w1118) and flies with specific ImpL2 overexpression (ImpL2^CDS^) at 24 hpi. tGPH, green; DAPI, cyan. The scale bar represents 20 µm.

ImpL2 is known to antagonize insulin signaling by binding to *Drosophila* insulin-like peptides (Honegger et al. 2008), which is here supported by effects on Foxo in the fat body, a known target of insulin signaling. Therefore, we tried to check the state of insulin signaling in the fat body by the commonly used PI3K reporter tGPH and phosphorylation of Akt. In response to infection, the tGPH reporter showed increased cytosolic localization in comparison to control flies when analyzed by confocal microscopy (Figure 5B). This effect was phenocopied by overexpression of ImpL2 in macrophages even in the absence of infection (Figure 5B), demonstrating the ability of macrophage-derived ImpL2 to antagonize insulin signaling in the fat body. Unfortunatelly, the effect of infection on tGPH localization was not strong enough to clearly see a difference after knocking down Impl2 in a double-blind evaluation. pAkt also appeared to be too variable during *S. pneumoniae* infection, precluding the reasonable use of Akt phosphorylation in our model.

### Macrophage-derived ImpL2 increases lipids and carbohydrates both in circulation and in macrophages

Stimulated glycogen and triglyceride catabolism is manifested by hyperglycemia and hyperlipidemia in the circulation of infected flies. Elevated titers of glucose, trehalose, glycerides and free fatty acids were detected in the hemolymph of these flies (Figure 6A and Figure6-figure supplement 1).

**Figure 6.**
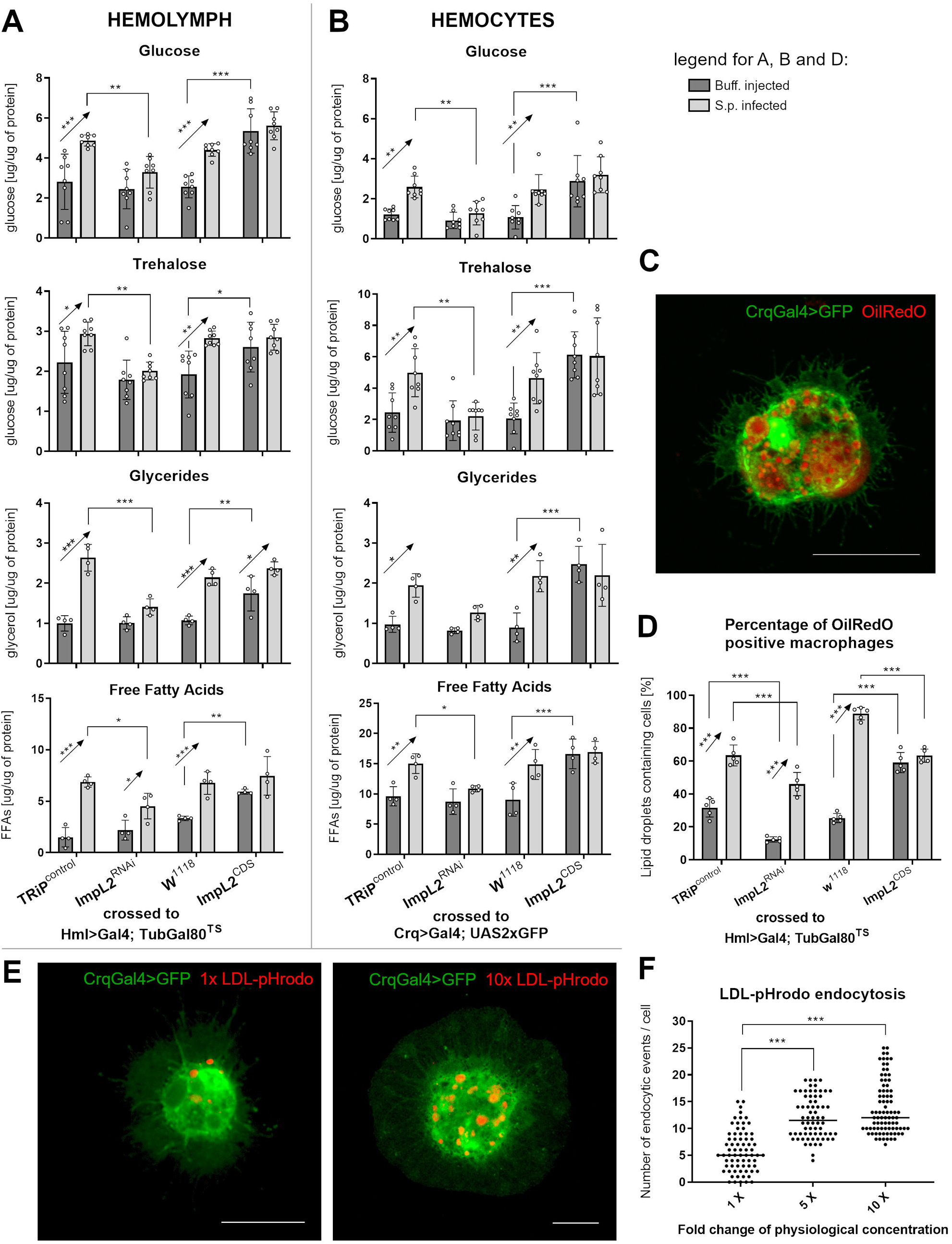
Macrophage-derived ImpL2 increases circulating carbohydrates and lipids to be available for activated macrophages. (**A**) Concentrations of circulating glucose, trehalose, glycerides, and free fatty acids in the hemolymph of Buff. injected and S.p. infected flies with macrophage-specific ImpL2 knockdown (ImpL2^RNAi^), overexpression (ImpL2^CDS^) and their respective controls (TRiP^control^, w1118) at 24 hpi. (**B**) Concentrations of glucose, trehalose, glycerides, and free fatty acids (**H**) in macrophages of Buff. injected and S.p. infected flies with macrophage-specific ImpL2 knockdown (ImpL2^RNAi^), overexpression (ImpL2^CDS^), and their respective controls (TRiP^control^, w1118) at 24 hpi. (**C**) Representative confocal microscopy image of macrophages (Crq>GFP) containing lipid droplets; neutral lipids were stained with OilRedO (red). The scale bar represents 10 µm. The image represents a Z-stack consisting of a maximum projection of 5 layers. (**D**) Proportional occurrence of macrophages containing at least one lipid droplet (stained by OilRedO) at 24 hpi, data combined from fifty Crq>GFP positive cells. (**E**) Representative confocal microscopy images depicting the ability of macrophage (Crq>GFP) to endocytose lipoproteins (human LDL-pHrodo, red). The scale bar represents 10 µm. (**F**) Number of endocytic events in macrophages of flies, which were injected with various concentrations (1x – dose corresponding to physiological concentration, 5x, and 10x) of lipoproteins (human LDL-pHrodo). The results are combined from three biological replicates. (A-B) Metabolite concentrations were normalized to the amount of proteins in each sample. Carbohydrate, resp. lipid measurements were performed in eight, resp. four replicates represented by individual dots. Results in (A,B and D) were compared by 2way ANOVA Tukey’s multiple comparisons test and in (**F**) one-way ANOVA Tukey’s multiple comparisons test. Bars/lines show means ± SD, asterisks mark statistically significant differences (*p<0.05; **p<0.01; ***p<0.001).

Increased levels of circulating lipids and carbohydrates were accompanied by increased accumulation of these energy-rich compounds in infection-activated macrophages (Figure 6B). These results are supported by the higher occurrence of lipid droplets in the cytosol of infection-activated macrophages after staining with the neutral lipid dye OilRedO (Figure 6C and D). The ability of macrophages to uptake mobilized lipids was verified by injection of fluorescently labeled lipoproteins (LDL-pHrodo; Figure 6E and F). These lipoproteins can be endocytosed via recognition by the scavenger receptor Croquemort (fly homolog of mammalian CD36), which is abundantly expressed in *Drosophila* macrophages. Injection of different concentrations of LDL-pHrodo showed that lipoprotein uptake by macrophages correlates with the amount of lipoproteins in the circulation, even beyond the physiological concentrations commonly occuring in the hemolymph (Gilbert and Chino 1974). Reserve mobilization, which leads to an increase in circulating nutrients and their subsequent accumulation in activated macrophages, depends on the production of ImpL2 by macrophages. Indeed, knockdown of macrophage-specific ImpL2 suppresses all observed effects stimulated by infection, whereas overexpression of ImpL2 mimicked these effects even in uninfected individuals (Figure 6A-D).

Taken together, the above results suggest that ImpL2 produced by macrophages during infection affects insulin signaling in adipose tissue, thereby triggering a Foxo-mediated transcriptional program that provides macrophages with the nutrients they require.

### ImpL2 is required for an effective immune response but can be detrimental during chronic infection

The efficiency of the immune response depends on an adequate supply of energy and essential precursors to activated immune cells. Therefore, we decided to investigate whether ImpL2-mediated release of reserves is necessary for an effective immune response. Lack of ImpL2 production by macrophages significantly reduced survival of flies infected with *S. pneumoniae* (Figure 7A and Figure 7-figure supplement 1). This infection is associated with elevated pathogen load (Figure 7B), indicating decreased resistance in these individuals. Elimination of bacteria and survival of *S. pneumonia* infection is critically dependent on efficient phagocytosis, otherwise flies succumb to infection within two days (Bajgar and Dolezal 2018). Indeed, reduced resistance in flies with knocked-down ImpL2 is associated with reduced phagocytic rates (Figure 7E and F). Infection-induced increases in antimicrobial peptide production by macrophages were reduced by ImpL2 knockdown for two of the three peptides analyzed (Figure 7-figure supplement 2). On the other hand, overexpression of ImpL2 24 hours prior to infection improved resistance to streptococcal infection, as evidenced by lower pathogen load and improved survival (Figure 6C and D). However, we did not detect a difference in phagocytosis and expression of the two antimicrobial peptides during infection is rather reduced in flies overexpressing ImpL2 (Figure 7-figure supplement 2), leaving the reasons for improved survival unclear.

**Figure 7.**
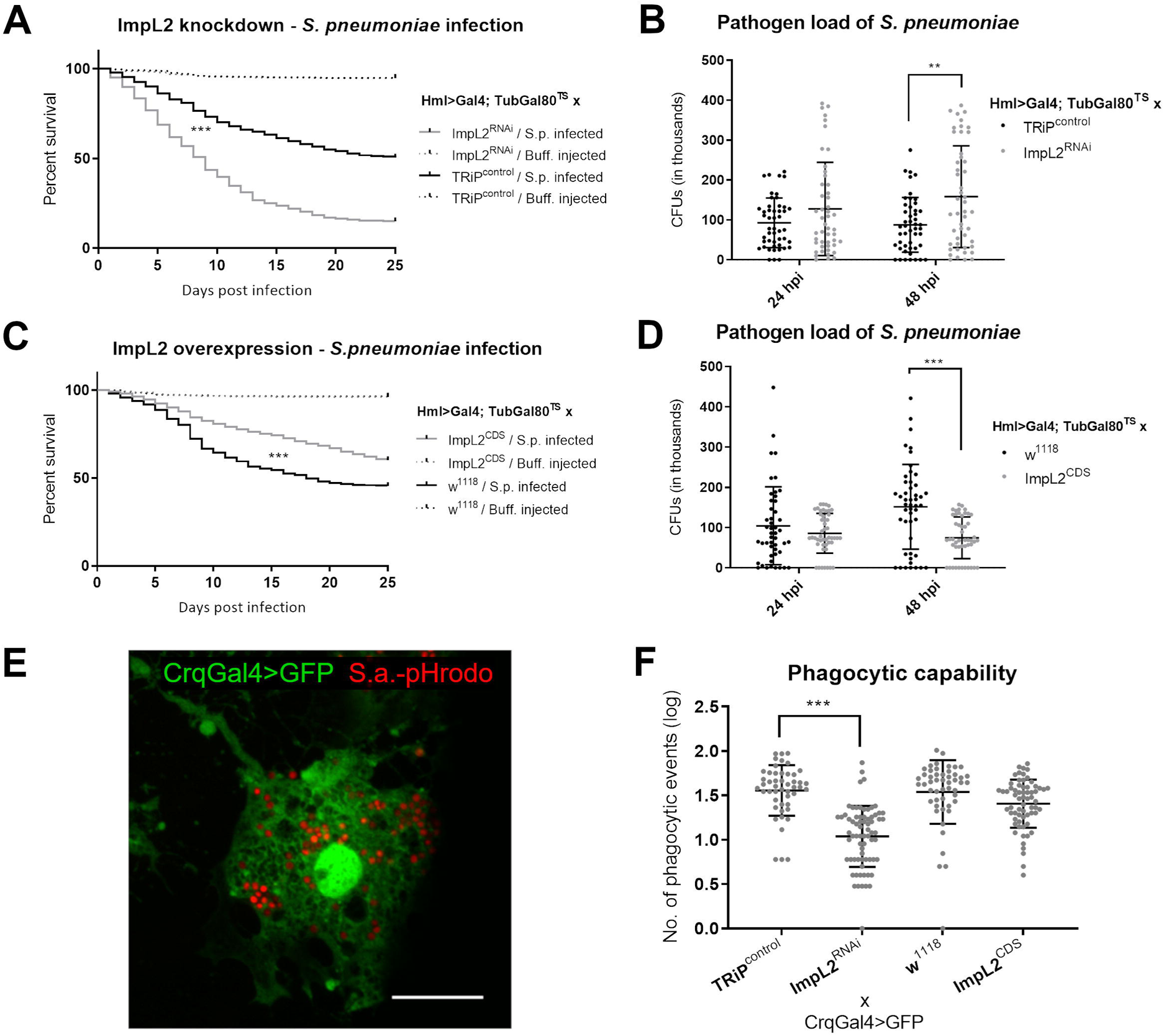
Macrophage-derived ImpL2 is important for resistance to bacterial infection. (**A, C**) Survival rate of S. pneumoniae (S.p.) infected and Buff. injected flies with macrophage-specific ImpL2 knockdown (ImpL2^RNAi^) (**A**), overexpression (ImpL2^CDS^) (**C**) and their respective controls (TRiP^control^, w1118). Three independent experiments were performed and combined into one survival curve; the number of individuals per replicate was at least 500 for each genotype. (**B, D**) Pathogen load in S.p. infected and Buff. injected flies with knockdown (ImpL2^RNAi^) (**B**), overexpression (ImpL2^CDS^) (**D**) and their respective controls (TRiP^control^, w1118) at 24 and 48 hpi. The individual dots in the plot represent the number of bacteria (colony forming units-CFUs) in one fly. The data show results combined from three independent biological replicates. (**E**) Confocal microscopy image depicting the ability of a macrophage (Crq>GFP) to phagocytose an invading pathogen (S. aureus-pHrodo-red). The scale bar represents 10 µm. The image represents a Z-stack consisting of a maximum projection of 5 layers. (**F**) Phagocytic rate calculated for flies with macrophage-specific ImpL2 knockdown (ImpL2^RNAi^), overexpression (ImpL2^CDS^), and their respective controls (TRiP^control^, w1118); each dot in the plot represents the log10-transformed number of phagocytic events per macrophage (Crq>GFP); results are combined from three independent experiments. Survival data (A and C) were analyzed by Log-rank and Grehan-Breslow Wilcoxon tests, pathogen load (B and D) by Mann-Whitney test and phagocytic capability (**F**) by one-way ANOVA Tukey’s multiple comparisons test. Lines are means ± SD, asterisks mark statistically significant differences (*p<0.05; **p<0.01; ***p<0.001).

Although our results demonstrate a beneficial role of ImpL2 during the acute phase of infection, ImpL2 has previously been associated with detrimental effects via induction of chronic cachexia-like wasting in a model of neoplastic tumor in *Drosophila* (Kwon et al. 2015). This suggests that the beneficial role of ImpL2 may be restricted to the short period of the acute phase of the immune response. Therefore, we tested the effects of ImpL2 manipulations in flies challenged with chronic infection caused by the intracellular pathogen *Listeria monocytogenes*. Flies injected with Listeria are unable to eliminate these bacteria and the length of their survival is determined by disease tolerance (Louie et al. 2016) whereas virulence and intracellular growth of *Listeria* depend on the availability of nutrients in the cytosol of the host cell (Chen, Pensinger, and Sauer 2017). Concordantly, the overexpression of ImpL2, which led to increased nutrient supplementation in immune cells, resulted in chronically increased *Listeria* burden (Figure 8A and B). This indicates that the originally beneficial effects of ImpL2 observed during the acute response to streptococcal infection may become detrimental during chronic immune challenge. Consistent with this, silencing of ImpL2 production by macrophage significantly reduced the intracellular load of *L. monocytogenes* at both 24 hpi and 12 days post-infection (Figure 8C and D), which may be explained by either a decreased ability of macrophages to engulf *Listeria* or a reduction of nutrients in their cytosol.

**Figure 8.**
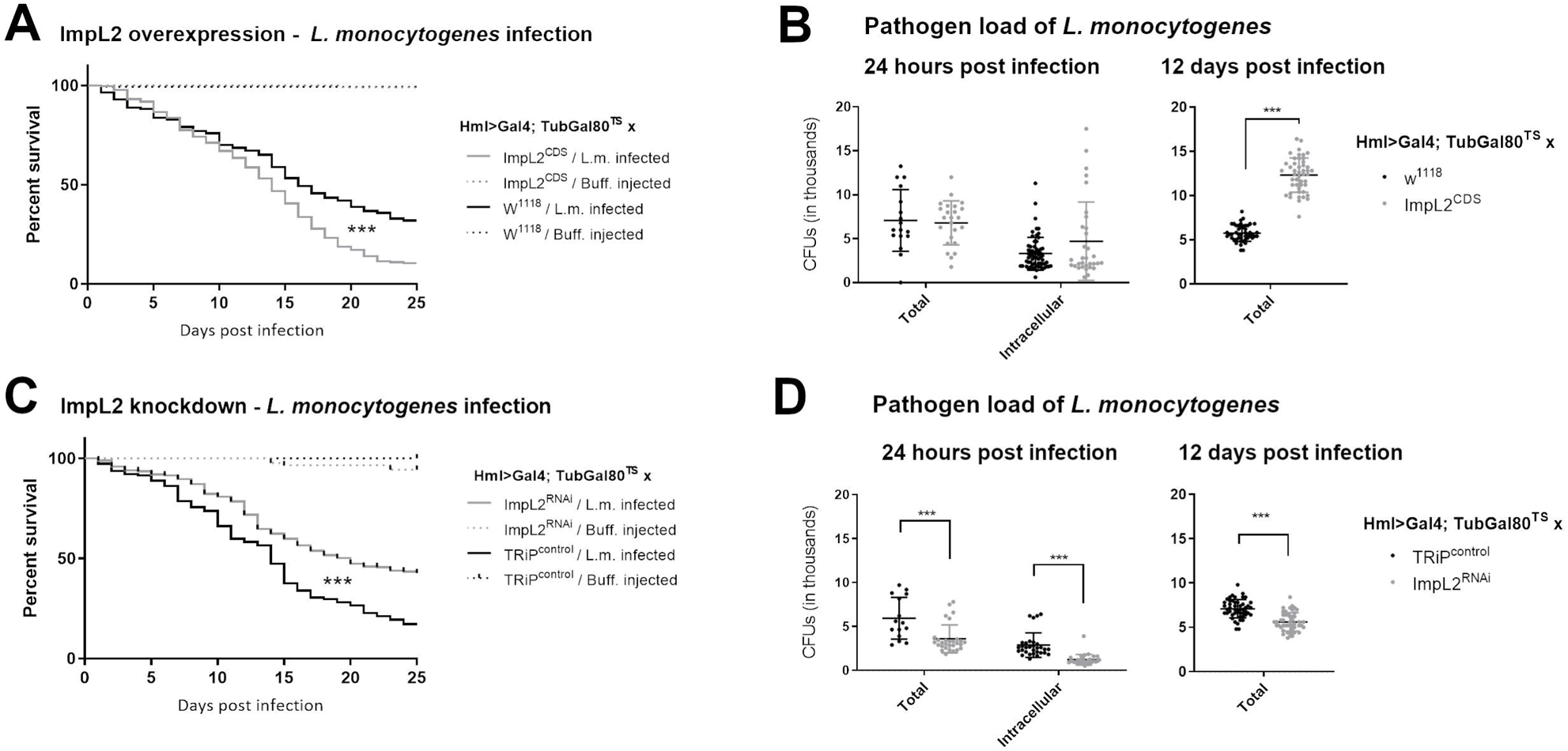
ImpL2 enhances deleterious effects of chronic infection. (**A, C**) Survival rate of L. monocytogenes (L.m.) infected and Buff. injected flies with macrophage-specific ImpL2 overexpression (ImpL2^CDS^) (**A**), knockdown (ImpL2^RNAi^) (**C**), and their respective controls (TRiP^control^, w1118). Three independent experiments were performed and combined into each survival curve; the number of individuals per replicate was at least 500 for each genotype. (**B, D**) Pathogen load of L. monocytogenes in flies with macrophage-specific ImpL2 overexpression (ImpL2^CDS^) (**B**), knockdown (ImpL2^RNAi^) (**D**), (**E**) and their respective controls (TRiP^control^, w1118) shown as either total load or intracellular Listeria subpopulation at 24 hpi and at 12 days post-infection. The individual dots in the plot represent the number of bacteria (colony forming units-CFUs) in one fly. The data show results combined from three independent biological replicates. Survival data (A and C) were analyzed by Log-rank and Grehan-Breslow Wilcoxon tests and pathogen load (B and D) by Mann-Whitney test. Values are mean ± SD, asterisks mark statistically significant differences (*p<0.05; **p<0.01; ***p<0.001).

In conclusion, ImpL2-mediated resource mobilization is essential for an adequate antibacterial immune response and resistance to infection. On the other hand, chronic or excessive production of ImpL2 tends to intensify the deleterious effects of chronic intracellular infections.

### Expression of IGFBP7, a mammalian homolog of ImpL2, is upregulated in immune-activated THP-1 *cells*

It has previously been shown that IGFBP7, the mammalian homologue of ImpL2, is produced by lipid-loaded liver macrophages in obese mice and human patients, with consequent effects on systemic metabolism through the regulation of insulin signaling in hepatocytes. ImpL2 released from lipid-loaded macrophages appears to play an analogous role in *Drosophila* fed a haigh-fat diet (Morgantini et al. 2019). To test the potential importance of the role of IGFBP7 in humans during infection, we measured the expression of IGFBPs in the PMA-activated THP-1 human monocytic cell line 24 hours after exposure to *S. pneumoniae* bacteria (Figure 9A). Expression of several IGFBP family members, namely *IGFBP1*, *IGFBP2*, *IGFBP5,* and *IGFBP6,* was not detected, indicating that these genes are not abundantly expressed in these cells (data not shown). Interestingly, *IGFBP7* gene expression was significantly increased in inoculated culture compared to PBS-treated controls (Figure 9B). In addition, the expression of the *IGFBP3* gene, which inhibits insulin resistance (Mohanraj et al. 2013), was reduced fourfold in activated THP-1 cells (Figure 9B), which is in agreement with the expected pattern associated with the adoption of bactericidal polarization by mammalian macrophages. This data therefore suggest that the increase in ImpL2 / IGFBP7 production by macrophages in response to infection is evolutionarily conserved in mammals.

**Figure 9.**
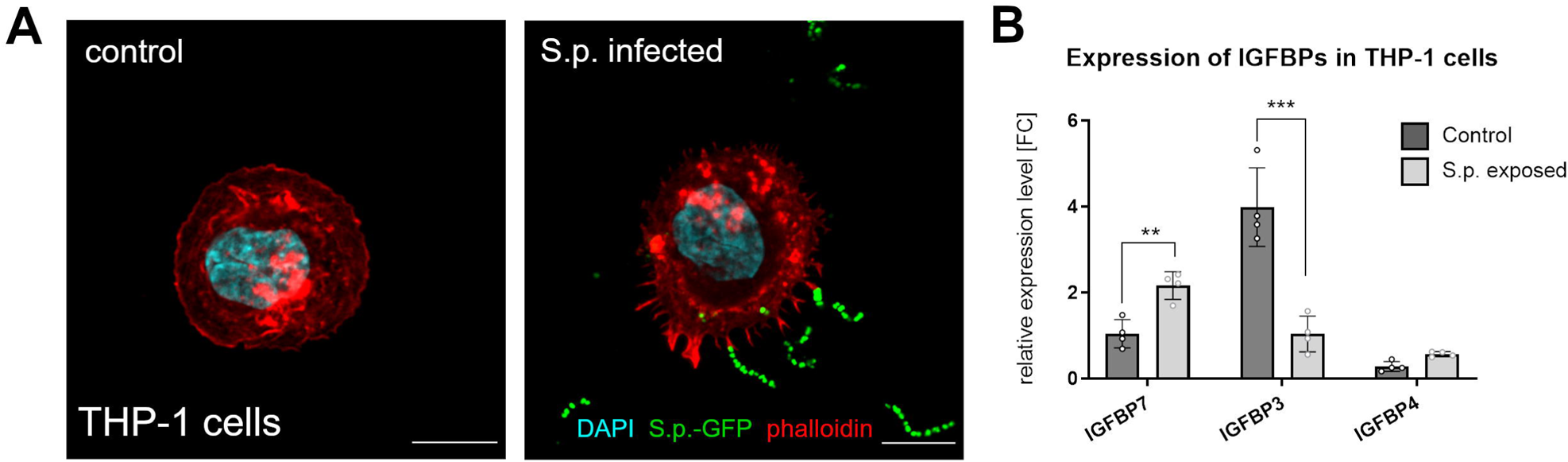
IGFBP7, the mammalian homolog of ImpL2, is produced in human macrophages in response to S. pneumoniae in vitro. (**A**) Confocal microscopy images of PMA-activated control THP-1 cells (left) and PMA-activated THP-1 cells actively interacting with GFP-labeled S. pneumoniae (green) in vitro (right). The scale bar represents 10 µm. The image on the right is a maximum projection of 3 layers. (**B**) Gene expression of IGFBP7, IGFBP3 and IGFBP4 in PMA-activated THP-1 cells 24 hours after exposure to S. pneumoniae in vitro. Expression levels normalized against ACTB are shown as a fold change relative to levels of IGFBP7 in controls that were arbitrarily set to 1. Measured data were compared in Graphpad Prism using 2way ANOVA Sidak’s multiple comparison test. Values are mean ± SD, asterisks mark statistically significant differences (*p<0.05; **p<0.01; ***p<0.001).

**Figure 9.**
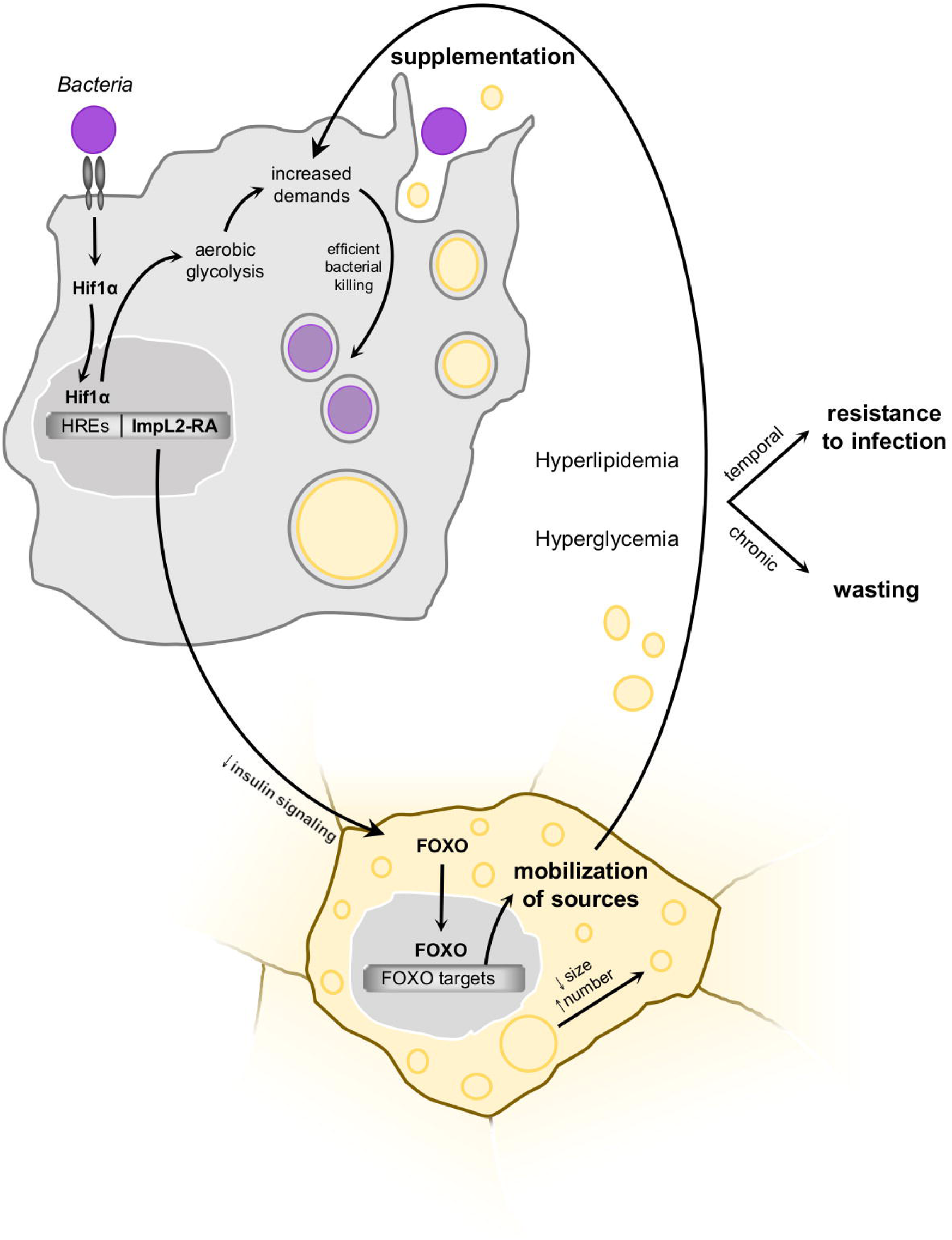
Schematic representation of the proposed model of ImpL2 function during infection. Upon recognition of the bacteria, the macrophages switch to Hif1α-driven aerobic glycolysis. Hif1α also induces the expression of the ImpL2-RA isoform by binding to hypoxia response elements in the promotor of the ImpL2 gene. Macrophage-derived ImpL2 then remotely activates Foxo re-localization to the nucleus in the fat body, leading to the expression of metabolic Foxo-target genes. Mobilized sources, manifested by hyperlipidemia and hyperglycemia in the hemolymph, are subsequently used by activated macrophages. These transient changes in ImpL2-regulated metabolism are essential for efficient bacterial killing and resistance to infection. However, if prolonged, for example during chronic infection, macrophage-derived ImpL2 can harm the host by potential wasting. Hif1α, hypoxia-inducible factor 1 α; HREs, hypoxia response elements; ImpL2, imaginal morphogenesis protein-late 2; Foxo, forkhead box O.

## Discussion

In this work, we show that infection-induced Hif1α transcriptional activity connects an intracellular metabolic switch to ImpL2 production in activated macrophages (Figre 10). ImpL2 subsequently induces the mobilization of resources from adipose tissue into the circulation, which may then become available to activated immune cells. This metabolic rearrangement of adipose tissue is most likely mediated by suppression of insulin signaling and subsequent transcriptional activity of Foxo. Such a program results in accelerated lipolysis and carbohydrate mobilization and the increased titer of circulating lipids and carbohydrates facilitates their utilization by macrophages. Although not directly tested in this work, these macromolecules are known to be important for bactericidal function of macrophages (Remmerie and Scott 2018). Indeed, here we show that such ImpL2-induced metabolic adaptation is essential for the acute immune response, but becomes maladaptive in the case of infection by intracellular pathogens that metabolically exploit the host cell. This is a remarkable example of how a beneficial metabolic program can become maladaptive in chronic diseases. This mechanism may not be limited to insects, indicated by an experiment showing that human macrophages activated by the same bacteria increase the expression of IGFBP7, the human homolog of *Drosophila* ImpL2.

We have previously shown that *Drosophila* macrophages undergo an evolutionarily conserved polarization to the M1 phenotype and thus their metabolism adjusts to Hif1α-induced aerobic glycolysis in response to bacterial challenge. This cellular metabolic shift is accompanied by systemic metabolic changes necessary to supplement the sudden nutritional needs of macrophages (Krejčová et al. 2019). Coordination of cellular metabolism in macrophages with systemic metabolism is crucial for resistance to infection and implies the existence of circulating factors mediating this interorgan crosstalk (Dolezal et al. 2019). Inspired by research on neoplastic cancer cells (Kwon et al. 2015; Figueroa-Clarevega and Bilder 2015; Bunker et al. 2015), in this study we focus on the cachectic factor ImpL2 as a candidate that has previously been linked to Hif1α transcriptional activity while having the potential to regulate systemic metabolism (Alee 2011; Li et al. 2013). To test whether the link between Hif1α activity and ImpL2 expression during hypoxia also applies to infection-activated macrophages, we performed a genomic *in silico* meta-analysis of hypoxic enhancer activity and Hif1α binding sites followed by Chip-qPCR to confirm the binding of Hif1α to previously identified genomic loci (Kamps-Hughes et al. 2015). This approach revealed the induction of a hypoxic enhancer in close proximity to the transcription start site of the *ImpL2-RA* transcript variant, which is further supported by the presence of four hypoxia response elements at this locus. *ImpL2-RA* is the most strongly expressed transcriptional variant in macrophages and its expression is further increased after infection. This infection-induced increase is Hif1α dependent, indicating a connection between ImpL2 expression and an internal metabolic switch in infection-activated macrophages. We have previously identified extracellular adenosine as a systemic factor derived from immune cells, with an analogous function to ImpL2, mobilizing carbohydrate stores to be available to the immune system (Bajgar et al. 2015; Bajgar and Dolezal 2018). While adenosine is a signaling molecule whose generation depends on the rate of cellular metabolism, the production of ImpL2 protein is linked to the transcriptional activity of the central metabolic regulator Hif1α. Thus, it is noteworthy that regulation of systemic metabolism can be mediated by factors with different mechanisms of production that reflect cellular metabolism of immune cells.

In addition to the regulation of carbohydrate metabolism, a prominent effect of ImpL2 is its regulation of systemic lipid metabolism. The metabolic changes observed in infection-challenged flies resemble lipemia as a characteristic symptom of sepsis in severely ill patients (Harris, Gosnell, and Kumwenda 2000). In sepsis-induced lipemia, insulin-resistant hepatocytes elevate the level of circulating lipids, which are preferentially utilized by activated immune cells in the periphery (Khovidhunkit et al. 2004; Aspichueta et al. 2012). Scavenger receptor-mediated endocytosis of serum lipids together with attenuated reverse cholesterol transport contribute substantially to cholesterol accumulation in bactericidal immune cells (Podrez et al. 2000). A number of immune-related macrophage functions depend on sufficient delivery of lipids, ranging from phagocytosis and phagolysosome maturation, to catecholamine production and immune memory formation via the mevalonate pathway (summarized in Remmerie and Scott 2018). Although providing macrophages with substantial amounts of lipids is essential for their proper function during the acute phase of infection, chronic exposure of macrophages to excessive lipids can lead to the adoption of a foam-cell phenotype, which promotes metabolic syndrome and atherosclerosis (Chistiakov et al. 2017; Febbraio, Guy, and Silverstein 2004).

Although we have not yet fully elucidate the mechanism of infection-induced lipid catabolism, we found that nuclear translocation of Foxo, followed by increased expression of several Foxo-target genes involved in this process, is regulated by macrophage-derived ImpL2. Foxo-driven mobilization of lipid stores requires suppression of insulin signaling in adipose tissue (Molaei, Vandehoef, and Karpac 2019; Luong et al. 2006), which is in agreemnet with the well-documented ability of ImpL2 to bind to *Drosophila* insulin-like peptides (DILPs) and thereby reduce insulin signaling in adipose tissue. This is further supported by our observations of reduced adipocyte insulin signaling in infected flies and flies with experimentally enhanced ImpL2 production in macrophages (Alee 2011; Alic et al. 2011; Arquier et al. 2006; Figueroa-Clarevega and Bilder 2015; Honegger et al. 2008; Kwon et al. 2015; Okamoto et al. 2013). Metabolic adjustments and the switch from anabolism to catabolism in adipose tissue during infection are regulated by multiple mechanisms (reviewed in Dolezal et al. 2019). Specifically, Foxo has previously been linked to wasting in flies during chronic infection (Dionne et al. 2006), suggesting its pathological effects during a prolonged immune response. In contrast, the remote effect of ImpL2 on reserve mobilization described in this work represents a rather beneficial role for ImpL2 during the acute immune response. Suppression of insulin signaling is often associated with chronic inflammation and can become detrimental to the organism by disrupting metabolic balance. However, the concept of selfish immunity (Straub 2014) considers insulin resistance as an evolutionary adaptive mechanism for rerouting nutrients toward the immune system during the acute immune response. Our results showing that ImpL2 is required for an effective immune response through changes in lipid and carbohydrate metabolism can be considered as experimental evidence for insulin resistance as an adaptive mechanism for resource mobilization in the acute phase of the immune response.

Neoplastic tumors share a characteristic cellular metabolism (Warburg 1925) with bactericidal macrophages (Andrejeva and Rathmell 2017). We suggest that they also share the production of ImpL2, which was originally perceived as a cancer-derived cachectic factor (Kwon et al. 2015; Figueroa-Clarevega and Bilder 2015), released from these cells to usurp energy from other tissues via cytokine-induced insulin resistance. The perception of ImpL2 expression as a consequence of cellular metabolic settings is supported by other conditions when ATP generation is independent of mitochondria, such as experimentally induced mitochondrial dysfunction, hypoxia, and neoplastic tumor growth (Alee 2011; Figueroa-Clarevega and Bilder 2015; Kwon et al. 2015; Li et al. 2013). Although this mechanism of metabolic regulation is beneficial to the organism during acute infection, it becomes detrimental in the case of tumor growth or chronic infection. Elimination of *S. pneumonia* requires effective phagocytosis – the flies either clear the infection or die. When phagocytosis is blocked, flies succumb to infection within two days (Bajgar and Dolezal 2018), so this immune response can be regarded as acute. ImpL2-mediated metabolic changes appear to be important for such a response. On the other hand, the harmful effect of ImpL2 on the survival of infection caused by the intracellular pathogen *L. monocytogenes* is demonstrated here. The flies are unable to eliminate these bacteria, which escape from the phagosome and establish a chronic intracellular infection from which the flies sooner or later die. In this case, reducing phagocytosis by knocking down ImpL2 lowered the short- and long-term intracellular load of the pathogen, leading to longer host survival. Various intracellular bacteria are known to take advantage of host cell supply and hijack their metabolic cascades to literarily nourish themselves (Péan et al. 2017; Teng, Ang, and Guan 2017). We do not know whether the observed increased *L. monocytogenes* burden in flies overexpressing ImpL2 is due to increased nutrients available to the pathogen, but these flies die more rapidly, indicating a rather negative role for ImpL2 during chronic infection.

In this study, we revealed that human macrophages activated by *S. pneumoniae* upregulate the expression of IGFBP7, the human homolog of *Drosophila* ImpL2, suggesting that a similar mechanism of metabolic regulation may operate in humans. Remarkably, not only infection but also experimentally induced metabolic syndrome in both *Drosophila* and mammals led to increased ImpL2/IGFBP7 expression in macrophages (Morgantini et al. 2019). While ImpL2-producing macrophages affect metabolism in the *Drosophila* fat body, which integrates functions of both mammalian adipose tissue and liver, IGFBP7-producing Kupffer cells act directly on hepatocytes to induce systemic metabolic changes (Morgantini et al. 2019; Akiel et al. 2017). The ability of both ImpL2 and IGFBP7 to bind extracellular insulin, thereby reducing systemic insulin signaling, further documents their functional homology, suggesting that their mechanism of action may also be conserved (Honegger et al. 2008; Arquier et al. 2006; Oh et al. 1996). Although this mechanism of resource mobilization may be important for the acute phase response, when prolonged it contributes to the progression of metabolic syndrome and atherosclerosis in mammals (Tomkin 2012). It is therefore not surprising that circulating IGFBP7 levels are considered a marker of systemic metabolic imbalance accompanying several human diseases such as obesity, acute kidney injury, liver fibrosis, and chronic obstructive pulmonary disease (Liu et al. 2015; Gunnerson et al. 2016; Martínez-Castillo et al. 2020; Ruan et al. 2017).

Combining our data with previous work, we conclude that the insulin/IGF antagonist ImpL2, which is released from activated macrophages as a reflection of their metabolic polarization, mediates nutrient mobilization from adipose tissue by reducing insulin signaling to liberate sufficient resources for the activated immune system. Despite the fact that both ImpL2 and insulin resistance are mostly studied in the context of pathological conditions such as cancer-induced cachexia, obesity, and chronic inflammatory conditions, our data revealed that they may play a beneficial role in the acute phase of bacterial infection. The relevance of our model is further supported by the production of IGFBP7, a homologue of ImpL2, in response to streptococcal infection in human macrophages. This leads us to the hypothesis that analogous mechanisms may also apply to macrophage-induced nutrient mobilization from hepatocytes during the acute phase of the immune response; however, this relationship remains to be experimentally verified.

## Funding

Tomáš Doležal - Czech Science Foundation (Project 20-09103S) Adam Bajgar - Czech Science Foundation (Project 20-14030S) Gabriela Krejčová - USB Grant Agency (Project 050/2019/P)

## Acknowledgment

The authors acknowledge funding from the Grant Agency of the Czech Republic to TD (Project 20-09103S; www.gacr.cz) and to AB (Project 20-14030S; www.gacr.cz). GK was supported by USB Grant Agency (Project 050/2019/P). We thank to Lucie Hrádková for laboratory services, enthusiasm and support, Alena Krejčí-Bruce and Lenka Chodáková for critical comments and inspiring discussions, Pavel Branný and Linda Doubravová for help with the preparation of the S.p.-GFP strain and Hana Sehadová for help with scanning electron microscopy. We thank to Hugo Stocker for the ImpL2^RNAi^, ImpL2^CDS^, and ImpL2-RA-Gal4 fly lines, Gabor Juhasz for the Atg8amCherry fly, and Marc Dionne for Crq>GFP fly line. Other fly stocks were obtained from the Bloomington Center (Bloomington, IN) and the VDRC (Vienna, Austria). The *S. pneumoniae* and *L. monocytogenes* bacterial strains were obtained from Dr. David Schneider. We also thank Martin Moss and Petr Šimek for the lipidomics service, the Department of Medical Biology (USB) for allowing us to use the S3eBioRad sorter, Biology Centre CAS for allowing us to use a confocal microscope, and a laboratory equipped to maintain human tissue cultures. We are also grateful to developers of Fiji: an open-source platform for biological-image analysis (doi:10.1038/nmeth.2019)

## Materials and methods

### Drosophila melanogaster strains and culture

The flies were raised on a diet containing cornmeal (80 g/l), sucrose (50 g/l), yeast (40 g/l), agar (10 g/l), and 10%-methylparaben (16.7 mL/l) and maintained in a humidity-controlled environment with a natural 12 h/12 h light/dark cycle at 25°C. Flies carrying Gal80 protein were raised at 18°C and transferred to 29°C 24 h prior to infection in order to degrade temperature-sensitive Gal80. Prior to the experiments, flies were kept in plastic vials on a sucrose-free cornmeal diet (cornmeal 53.5 g/l, yeast 28.2 g/l, agar 6.2 g/l and 10%-methylparaben 16.7 mL/l) for 7 days. Flies infected with *S. pneumoniae* were kept on a sucrose-free cornmeal diet in incubators at 29°C due to the temperature sensitivity of *S. pneumoniae*. They were transferred to fresh vials every other day without the use of CO_2_ to ensure good food condition. Flies infected with *L. monocytogenes* were kept on a sucrose-free cornmeal diet at 25°C. The Drosophila Stock Center in Bloomington provided *ImpL2^RNAi^* (*y1 sc* v1; P{TRiP.HMC03863}attP40*; FBst0055855) *TRiP*^control^ (*y[1] v[1]; P{y[+t7.7]=CaryP}attP40*; FBst0036304) and *20xUAS-6xmCherry* (*P{20XUAS-6XmCherry-HA}attP2*; FBtp0094992) flies. *ImpL2-RA-Gal4* (FBal0290965) and *UAS-ImpL2*^cds^ (*UAS-s.ImpL2*; FBal0249386) were kind gifts from Hugo Stocker. *CrqGal4>2xeGFP* were obtained from Marc Dionne. The *Atg8a-mCherry* strain was kindly provided by Gabor Juhasz. The *w*^1118^ strain has a genetic background based on *CantonS*.

### Genotypes of experimental models

**Figure 1**

**ImpL2-RA>mCherry** corresponds to *w^1118^/w^1118^*; 20xUAS-6xmCherry/+; ImpL2-RA-Gal4/+

**HmlGal4>GFP** refers to *w^1118^/w^1118^*; HmlΔ-Gal4 UAS-2xeGFP/HmlΔ-Gal4 UAS-2xeGFP; +/+

**HmlGal4>GFP; Hif1α^RNAi^** corresponds to *w^1118^*/+; HmlΔ-Gal4 UAS-2xeGFP/+; UAS-Hif1α^RNAi^/+

**ImpL2^RNAi^** refers to *w^1118^*/+; HmlΔ-Gal4/UAS-ImpL2*^RNAi^*; P{tubPGal80ts}/+

**ImpL2^cds^** corresponds to *w^1118^/w^1118^*; HmlΔ-Gal4/+; P{tubPGal80ts}/UAS-ImpL2^cds^

**TRiP^control^** refers to *w^1118^*/+; HmlΔ-Gal4/+; P{tubPGal80ts}/TRiP^control^

**w^1118^** corresponds to *w^1118^/w^1118^*; HmlΔ-Gal4/+; P{tubPGal80ts}/+

**Figure 1-figure supplement 3**

**Hml>Gal4, UAS-GFP x TRiP^control^** corresponds to *w^1118^*/+; HmlΔ-Gal4 UAS-2xeGFP/+; TRiP^control^/+

**Hml>Gal4, UAS-GFP x Hif1α^RNAi^** corresponds to *w^1118^*/+; HmlΔ-Gal4 UAS-2xeGFP/+; UAS-Hif1α^RNAi^/+

**Figure 2**

**ImpL2^RNAi^** refers to *w^1118^*/+; HmlΔ-Gal4/UAS-ImpL2^RNAi^; P{tubPGal80ts}/+

**ImpL2^cds^** corresponds to *w^1118^/w^1118^*; HmlΔ-Gal4/+; P{tubPGal80ts}/UAS-ImpL2^cds^

**TRiP^control^** refers to *w^1118^*/+; HmlΔ-Gal4/+; P{tubPGal80ts}/TRiP^control^

**w^1118^** corresponds to *w^1118^/w^1118^*; HmlΔ-Gal4/+; P{tubPGal80ts}/+

**Figure 3**

**ImpL2^RNAi^** refers to *w^1118^*/+; HmlΔ-Gal4/UAS-ImpL2^RNAi^; P{tubPGal80ts}/+

**ImpL2^cds^** corresponds to *w^1118^/w^1118^*; HmlΔ-Gal4/+; P{tubPGal80ts}/UAS-ImpL2^cds^

**Atg8a-mCherry** refers to gen +/+; Atg8a-mCherry/Atg8a-mCherry

**Figure 3-figure supplement 1**

**ImpL2RNAi** refers to *w^1118^*/+; HmlΔ-Gal4/UAS-ImpL2^RNAi^; P{tubPGal80ts}/+

**ImpL2^cds^** corresponds to *w^1118^/w^1118^*; HmlΔ-Gal4/+; P{tubPGal80ts}/UAS-ImpL2^cds^

**TRiP^control^** refers to *w^1118^*/+; HmlΔ-Gal4/+; P{tubPGal80ts}/TRiP^control^

**W^1118^** corresponds to *w^1118^/w^1118^*; HmlΔ-Gal4/+; P{tubPGal80ts}/+

**Figure 4**

**ImpL2^RNAi^** refers to *w^1118^*/+; HmlΔ-Gal4/UAS-ImpL2^RNAi^; P{tubPGal80ts}/+

**ImpL2^cds^** corresponds to *w^1118^/w^1118^*; HmlΔ-Gal4/+; P{tubPGal80ts}/UAS-ImpL2^cds^

**TRiP^control^** refers to *w^1118^*/+; HmlΔ-Gal4/+; P{tubPGal80ts}/TRiP^control^

**w^1118^** corresponds to *w^1118^/w^1118^*; HmlΔ-Gal4/+; P{tubPGal80ts}/+

**Figure 5**

**Hml>Gal4 TubGal80 x w^1118^** corresponds to *w^1118^/w^1118^*; HmlΔ-Gal4 P{tubPGal80ts}/+; +/+

**Hml>Gal4 TubGal80 x ImpL2^cds^** corresponds to *w^1118^/w^1118^*; HmlΔ-Gal4 P{tubPGal80ts}/+; UAS-ImpL2^cd^/+

**Hml>Gal4 TubGal80 x foxo^BG01018^ ImpL2^cds^** refers to *w^1118^/w^1118^*; HmlΔ-Gal4 P{tubPGal80ts}/+; P{w[+mGT]=GT1}foxo^BG01018^ *UAS-ImpL2*^cds^/+

**Hml>Gal4 TubGal80; tGPH-GFP x w^1118^** corresponds to *w^1118^/w^1118^*; HmlΔ-Gal4 P{tubPGal80ts}/+; tGPH/+

**Hml>Gal4 TubGal80; tGPH-GFP x ImpL2^cds^** corresponds to *w^1118^/w^1118^*; HmlΔ-Gal4 P{tubPGal80ts}/+; tGPH/UAS-ImpL2^cds^

**Figure 6**

**ImpL2^RNAi^** refers to *w^1118^*/+; HmlΔ-Gal4/UAS-ImpL2^RNAi^; P{tubPGal80ts}/+

**ImpL2^cds^** corresponds to *w^1118^/w^1118^*; HmlΔ-Gal4/+; P{tubPGal80ts}/UAS-ImpL2^cds^

**TRiP^control^** refers to *w^1118^*/+; HmlΔ-Gal4/+; P{tubPGal80ts}/TRiP^control^

**w^1118^** corresponds to *w^1118^/w^1118^*; HmlΔ-Gal4/+; P{tubPGal80ts}/+

**Crq>Gal4; UAS2xGFP x ImpL2^RNAi^** refers to *w^1118^*/+; UAS-ImpL2^RNAi^/+; Crq-Gal4, UAS-2xeGFP/+

**Crq>Gal4; UAS2xGFP x ImpL2^cds^** corresponds to *w^1118^/w^1118^*; +/+; Crq-Gal4, UAS-2xeGFP/UAS-ImpL2^cds^

**Crq>Gal4; UAS2xGFP x TRiP^control^** refers to *w^1118^*/+; +/+; Crq-Gal4, UAS-2xeGFP/TRiP^control^

**Crq>Gal4; UAS2xGFP x W^1118^** corresponds to *w^1118^/w^1118^*; Crq-Gal4, UAS-2xeGFP/+

**CrqGal4>GFP** refers to *w^1118^/w^1118^*; Crq-Gal4, UAS-2xeGFP/Crq-Gal4, UAS-2xeGFP

**Figure 7**

**ImpL2^RNAi^** refers to *w^1118^*/+; HmlΔ-Gal4/UAS-ImpL2^RNAi^; P{tubPGal80ts}/+

**ImpL2^cds^** corresponds to *w^1118^/w^1118^*; HmlΔ-Gal4/+; P{tubPGal80ts}/UAS-ImpL2^cds^

**TRiP^control^** refers to *w^1118^*/+; HmlΔ-Gal4/+; P{tubPGal80ts}/TRiP^control^

**w^1118^** corresponds to *w^1118^/w^1118^*; HmlΔ-Gal4/+; P{tubPGal80ts}/+

**Crq>Gal4 x ImpL2^cds^** corresponds to *w^1118^/w^1118^*; +/+; Crq-Gal4, UAS-2xeGFP/UAS-ImpL2^cds^

**Figure 7-figure supplement 1**

**Hml-Gal4; TubGal80^TS^** refers to *w^1118^/w^1118^*; HmlΔ-Gal4/+; P{tubPGal80ts}/+

**Hml-Gal4; TubGal80^TS^ > ImpL2^cds^** corresponds to *w^1118^/w^1118^*; HmlΔ-Gal4/+; P{tubPGal80ts}/UAS-ImpL2^cds^

**ImpL2^cds^ x w^1118^** corresponds to *w^1118^/w^1118^*; +/UAS-ImpL2^cds^

**Hml-Gal4; TubGal80^TS^ > ImpL2^RNAi^** corresponds to *w^1118^*/+; HmlΔ-Gal4/UAS-ImpL2^RNAi^; P{tubPGal80ts}/+

**ImpL2^RNAi^** corresponds to *w^1118^*/+; +/UAS-ImpL2^RNAi^

**Figure 7-figure supplement 2**

**ImpL2^RNAi^** refers to *w^1118^*/+; HmlΔ-Gal4/UAS-ImpL2^RNAi^; P{tubPGal80ts}/+

**ImpL2^cds^** corresponds to *w^1118^/w^1118^*; HmlΔ-Gal4/+; P{tubPGal80ts}/UAS-ImpL2^cds^

**TRiP^control^** refers to *w^1118^*/+; HmlΔ-Gal4/+; P{tubPGal80ts}/TRiP^control^

**w^1118^** corresponds to *w^1118^/w^1118^*; HmlΔ-Gal4/+; P{tubPGal80ts}/+

**Figure 8**

**ImpL2^RNAi^** refers to *w^1118^*/+; HmlΔ-Gal4/UAS-ImpL2^RNAi^; P{tubPGal80ts}/+

**ImpL2^cds^** corresponds to *w^1118^/w^1118^*; HmlΔ-Gal4/+; P{tubPGal80ts}/UAS-ImpL2^cds^

**TRiP^control^** refers to *w^1118^*/+; HmlΔ-Gal4/+; P{tubPGal80ts}/TRiP^control^

**w^1118^** corresponds to *w^1118^/w^1118^*; HmlΔ-Gal4/+; P{tubPGal80ts}/+

### Hypoxic enhancer activity analysis

Genome-wide experimental data characterizing hypoxia-induced transcriptional enhancer activity (Kamps-Hughes et al. 2015) was used to analyze the ImpL2 region for hypoxic enhancers. Hypoxic transcriptional induction is defined as the ratio of expressed randomer tag counts in hypoxic versus normoxic conditions and was binned by 500-bp regions across the *ImpL2* locus. Each 500-bp bin was then analyzed for transcription factor binding sites corresponding to hypoxic (Hif1α), immune (Rel) and stress (HSF) response transcription factors. Transcription factor position frequency matrices were downloaded from the JASPAR database (https://academic.oup.com/nar/article/32/suppl_1/D91/2505159) and queried against the 500-bp bin sequences using BOBRO software (https://www.ncbi.nlm.nih.gov/pmc/articles/PMC3074163/). BOBRO generates a p-value of enrichment for the position frequency matrix within the search sequence and also identifies the location of the predicted binding sites.

### Chip-qPCR assay

The Pro-A Drosophila CHIP Seq Kit (Chromatrap) was used to co-immunoprecipitate genomic regions specifically bound by the transcription factor HIF1α. A transgenic fly strain carrying the Hif1α protein fused to GFP (BDSC: 42672) was used for this purpose. The procedure was performed according to the supplier’s instructions. Briefly, the slurry was prepared by homogenizing ten infected or PBS injected males in three biological replicates. The Rabbit Anti-GFP antibody (ABfinity) was bound to the chromatographic column. Genomic DNA was fragmented to an approximate size of 500 bp by three cycles of 60-second sonication. The fragment size was verified by gel electrophoresis. All samples were tested with positive and negative controls. The amount of precipitated genomic fragments was normalized to the amount of fragments in slurry before precipitation. The ImpL2-RA promoter sequence was covered with seven primer pairs corresponding to the 500-bp bins upstream of the transcription start site previously assessed in the *in silico* analysis. The genomic region of S-adenosylmethionine synthetase was used as a negative control since it does not contain any sequences of hypoxia response elements. Primer sequences are listed in the Key Resources Table. The amount of HIF1α-bound regions of the ImpL2 promoter was quantified on a 96CFX 1000 Touch Real-Time Cycler (BioRad) using TP 2x SYBR Master Mix (Top-Bio) in three technical replicates with the following protocol: initial denaturation - 3 min at 95°C, amplification – 15 s at 94°C, 20 s at 56°C, 25 s at 72°C for 40 cycles. Melting curve analysis was performed at 65–85°C/step 0.5°C. The qPCR data were analyzed using double delta Ct analysis

### Gene expression analysis

Gene expression analyzes were performed on several types of samples: six whole flies, six thoraxes, six fat bodies, or 20,000 isolated macrophages. Macrophages were isolated by a cell sorter (S3e Cell Sorter, BioRad) as described in the section Isolation of macrophages, while dissections were made in PBS, transferred to TRIzol Reagent (Invitrogen) and homogenized using a DEPC-treated pestle. Subsequently, RNA was extracted by TRIzol Reagent (Invitrogen) according to the manufacturer’s protocol. Superscript III Reverse Transcriptase (Invitrogen) primed by oligo(dT)20 primer was used for reverse transcription. Relative expression rates for particular genes were quantified on a 384CFX 1000 Touch Real-Time Cycler (BioRad) using the TP 2x SYBR Master Mix (Top-Bio) in three technical replicates with the following protocol: initial denaturation - 3 min at 95°C, amplification – 15 s at 94°C, 20 s at 56°C, 25 s at 72°C for 40 cycles. Melting curve analysis was performed at 65–85°C/step 0.5°C. The primer sequences are listed in the Key Resources Table. The qPCR data were analyzed using double delta Ct analysis, and the expressions or specific genes were normalized to the expression of Ribosomal protein 49 (Rp49) in the corresponding sample. The relative values (fold change) to control are shown in the graphs. Samples for gene expression analysis were collected from three independent experiments.

### Metabolite measurement

To measure metabolite concentration, isolated macrophages, whole flies or hemolymph were used. Hemolymph was isolated from 25 adult males by centrifugation (14,000 RPM, 5 min) through a silicagel filter into 50 μL of PBS. For measurement of metabolites from whole flies, five flies were homogenized in 200 μL of PBS and centrifuged (3 min, 4°C, 8,000 RPM) to discard insoluble debris. 50,000 macrophages were isolated by cell sorter (S3e Cell Sorter, BioRad) as described in the section Isolation of macrophages. Half of all samples were used for the quantification of proteins. Samples for glucose, glycogen, trehalose, and triglyceride measurement were denatured at 75°C for 10 min, whereas samples for protein quantification were frozen immediately in −80°C. Glucose was measured using a Glucose (GO) Assay (GAGO-20) Kit (Sigma) according to the manufacturer’s protocol. Colorimetric reaction was measured at 540 nm. For glycogen quantification, sample was mixed with amyloglucosidase (Sigma) and incubated at 37°C for 30 min. A Bicinchoninic Acid Assay (BCA) Kit (Sigma) was used for protein quantification according to the supplier’s protocol and the absorbance was measured at 595 nm. Cholesterol and cholesteryl esters were measured on isolated lipid fraction by using Cholesterol/Cholesteryl Ester Quantitation Kit (Sigma) according to the supplieŕs protocol. Triglycerides were measured using Triglyceride quantification Colorimetric/Fluorometric Kit (Sigma). For trehalose quantification, sample was mixed with trehalase (Sigma) and incubated at 37°C for 30 min. Samples for metabolite concentration were collected from three independent experiments.

### Staining of lipid droplets

Flies were dissected in Gracés Insect Medium (Sigma) and subsequently stained with DAPI and Cell Brite Fix Membrane Stain 488 (Biotium) diluted with Gracés Insect Medium according to the manufactureŕs protocol at 37°C. Tissues were washed in PBS and then fixed with 4% PFA (Polysciences). After 20 min, the tissues were washed in PBS and pre-washed with 60% isopropanol. Dissected abdomens were then stained with OilRedO dissolved in 60% isopropanol for 10 min. The tissues were then washed with 60% isopropanol and mounted in an Aqua Polymount (Polysciences). Tissues were imaged using an Olympus FluoView 3000 confocal microscope (Olympus). Content of lipids in adipose tissue and size of lipid droplets were analyzed using Fiji software. Flies for the analysis of lipid droplets in the fat body were collected from three independent experiments and representative images are shown.

### Lipidomic analysis

Adipose tissues from six flies from each group were dissected in ice-cold PBS for lipidomics analysis. The removed tissues were stored in PBS buffer in Eppendorf tubes at – 80 °C. Immediately after thawing, they were extracted by 500 µl of cold chloroform: methanol solution (v/v; 1:1). The samples were then homogenized by a Tissue Lyser II (Qiagen, Prague, Czech Republic) at 50 Hz, −18°C for 5 min and kept further in an ultrasonic bath (0°C, 5 min). Further, the mixture was centrifuged at 10,000 RPM at 4°C for 10 min followed by the removal of the supernatant. The extraction step was repeated at the same conditions. The lower layer of pooled supernatant was evaporated to dryness under a gentle stream of Argon. The dry total lipid extract was re-dissolved in 500 µl of chloroform: methanol solution (v/v; 1:1) and directly measured using previously described methods (Bayley et al. 2020). Briefly, high performance liquid chromatography (Accela 600 pump, Accela AS autosampler) combined with mass spectrometry LTQ-XL (all Thermo Fisher Scientific, San Jose, CA, USA) were used. The chromatographic conditions were as follows: Injection volume 5 µl; column Gemini 3 µM C18 HPLC column (150 × 2 mm ID, Phenomenex, Torrance, CA, USA) at 35°C; the mobile phase (**A**) 5 mM ammonium acetate in methanol with ammonia (0.025%), (**B**) water and (**C**) isopropanol: MeOH (8:2); gradient change of A:B:C as follows: 0 min: 92:8:0, 7 min: 97:3:0, 12 min: 100:0:0, 19 min: 93:0:7, 20-29 min: 90:0:10, 40-45 min: 40:0:60, 48 min: 100:0:0, and 50-65 min: 92:8:0 with flow rate 200 µl/min. The mass spectrometry condition: positive (3 kV) and negative (−2,5 kV) ion detection mode; capillary temperature 200°C. Eluted ions were detected with full scan mode from 200 to 1000 Da with the collisionally induced MS2 fragmentation (NCE 35). Data were acquired and processed by means of XCalibur 4.0 software (Thermo Fisher). The corrected areas under individual analytical peaks were expressed in percentages assuming that the total area of all detected is 100%.

### Autophagy visualization

To visualize autophagy in adipose tissue, Atg8a-mCherry-bearing flies were dissected in PBS, fixed with 4% PFA, and then washed with PBS and stained with DAPI. Fat bodies were imaged using an Olympus FluoView 3000 confocal microscope (Olympus).

### Immunostaining

Flies were dissected in ice-cold PBS and fixed with 4% PFA in PBS (Polysciences) for 20 minutes. After three washes in PBS-Tween (0.1%), nonspecific binding was blocked by 10% NGS in PBS for 1 hour at RT. Tissues were then incubated with primary antibodis (for NimC1: Mouse anti-NimC1 antibody P1a+b, 1:100, kindly provided by István Andó); for Foxo: Rabbit anti-Foxo, CosmoBio, 1:1,000; for tGPH: Rabbit anti-GFP, ABfinity, 1:100) at 4°C overnight. After washing the unbound primary antibody (three times for 10 min in PBS-Tween), secondary antibody was applied at a dilution of 1:250 for 2 hours at RT (Goat anti-Mouse IgG (H+L) Alexa 555, Invitogen or Goat anti-Rabbit IgG (H+L) Cy2, Jackson-Immunoresearch). Nuclei were stained with DAPI. Tissues were mounted with Aqua Polymount (Polysciences). Tissues were imaged using an Olympus FluoView 3000 confocal microscope (Olympus) and images were reconstructed using Fiji software. Foxo localization was detected by Plot-Profile analysis using Fiji software.

### Isolation of macrophages

GFP-labeled macrophages were isolated from *Crq-Gal4, UAS-eGFP* male flies using fluorescence-activated cell sorting (FACS). Approximately 200 flies were anaesthetized with CO_2_, washed in PBS and homogenized in 600 mL of PBS using a pestle. The homogenate was sieved through a nylon cell strainer (40 μm). This strainer was then additionally washed with 200 μL of PBS, which was added to the homogenate subsequently. The samples were centrifuged (3 min, 4°C, 3,500 RPM) and the supernatant was washed with ice-cold PBS after each centrifugation (three times). Prior to sorting, samples were transferred to FACS polystyrene tubes using a disposable bacterial filter (50 μm, Sysmex) and macrophages were sorted into 100 μL of PBS using a S3^TM^ Cell Sorter (BioRad). Isolated cells were verified by fluorescence microscopy and differential interference contrast.

### Phagocytic activity

To analyze phagocytic rate, flies were infected with 20,000 of *S. pneumoniae* and after 24 h, they were injected with 50 nl of pHrodo^TM^ Red *S. aureus* (Thermo Fischer Scientific). After 1 h, abdomens of flies were dissected in PBS and then fixed for 20 min with 4% PFA. Aqua Polymount (Polysciences) was used to mount the sample. Macrophages were imaged using an Olympus FluoView 3000 confocal microscope and red dots depicting phagocytic events were manually counted per cell.

### Lipoprotein uptake

To analyze lipoprotein uptake by *Drosophila* macrophages, Crq>GFP flies were injected with 50 nl of 1x (corresponding to physiological concentration), 5x or 10x concentrated pHrodo^TM^ Red LDL (Invitrogen) into the ventrolateral side of the abdomen using an Eppendorf Femtojet microinjector. After 1 hour, the fly abdomens were opened in PBS and subsequently fixed for 20 min with 4% PFA in PBS (Polysciences). Aqua Polymount (Polysciences) was used to mount the sample. Macrophages were imaged using an Olympus FluoView 3000 confocal microscope.

### Bacterial strain and fly injection

The *Streptococcus pneumoniae* strain EJ1 was stored at −80°C in Tryptic Soy Broth (TSB) media containing 10% glycerol. For the experiments, bacteria were streaked onto agar plates containing 3% TSB and 100 mg/mL streptomycin and subsequently incubated at 37°C in 5% CO_2_ overnight. Single colonies were inoculated into 3 mL of TSB liquid media with 100 mg/mL of streptomycin and 100,000 units of catalase and incubated at 37°C + 5% CO2 overnight. Bacterial density was measured after an additional 4 h so that it reached an approximate 0.4 OD600. Final bacterial cultures were centrifuged and dissolved in PBS so the final OD reached 2.4. The *S. pneumoniae* culture was kept on ice prior to injection and during the injection itself. Seven-day-old males were anaesthetized with CO_2_ and injected with 50 nL culture containing 20,000 *S. pneumoniae* bacteria or 50 nL of mock buffer (PBS) into the ventrolateral side of the abdomen using an Eppendorf Femtojet microinjector. The *Listeria monocytogenes* strain 10403S was stored at −80°C in brain and heart infusion (BHI) broth containing 25% glycerol. For the experiments bacteria were streaked onto Luria Bertani (LB) agar plates containing 100 μg/mL streptomycin and incubated at 37°C overnight; plates were stored at 4°C and used for inoculation for a period of two weeks. Single colonies were used to inoculate 3 mL of BHI and incubated overnight at 37°C without shaking to obtain a morning 600 nm optical density (OD600) of approx. 0.4. Further, *L*. *monocytogenes* cultures were diluted to OD600 0.01 in phosphate buffered saline (PBS) and stored on ice prior to loading into an injection needle. Approximately 1,000 *Listeria* per fly were injected.

### Survival analysis

*Streptococcus*-injected flies were kept at 29°C in vials with approximately 30 individuals per vial and were transferred to fresh food every other day. *Listeria*-injected flies were kept at 25°C. Dead flies were counted daily. At least three independent experiments were performed and combined into a single survival curve generated in Graphpad Prism software; individual experiments showed comparable results. The average number of individuals was more than 500 for each genotype and replicate.

### Pathogen load measurement

Single flies were homogenized in PBS using a motorized plastic pestle in 1.5 mL tubes. Bacteria were plated in spots onto LB (*L*. *monocytogenes*) or TSB (*S*. *pneumoniae*) agar plates containing streptomycin in serial dilutions and incubated overnight at 37°C before manual counting. To determine intracellular *L*. *monocytogenes* loads, flies were injected with 50 nL of gentamycin solution (1 mg/mL in PBS) 3 h prior to fly homogenization. Pathogen loads of 16 flies were determined for each genotype and treatment in each experiment; at least three independent infection experiments were conducted and the results were combined into one graph (in all presented cases, individual experiments showed comparable results).

### THP-1 cell lines

THP-1 cells were cultured at 37°C, 5% CO_2_, in RPMI-1640 medium (Sigma), supplemented with 2 mM glutamine (Applichem), 2 g/L sodium bicarbonate (J&K Scientific), 10% FBS (Biosera) and 100 mg/L streptomycin (Sigma). Prior to the experiment, cells were transferred to 24-well plates at 10^5^ cells/well in four biological replicates. THP-1 cells were activated with phorbol-12-myristate-13-acetate (200 ng/mL, MedChemExpress). After 24 hours, *S. pneumoniae* bacteria were added (MOI 50 bacteria/macrophage) and the plate was centrifuged briefly (2 min, 200xg). Following 6-h incubation, the cells were washed with RPMI-1640 medium, and fresh RPMI-1640 supplemented with gentamycin (0.1 mg/mL, Sigma) was added. After 1 h incubation, the medium was replaced with RPMI-1640 supplemented with penicillin-streptomycin (1%, Biosera). After another 17 h, the cells were harvested into TRIzol Reagent (Invitrogen) followed by RNA isolation.

### Statistics

All data were analyzed using Graphpad Prism software; specific tests are listed for each graph in the figure legend.

### Key resources table

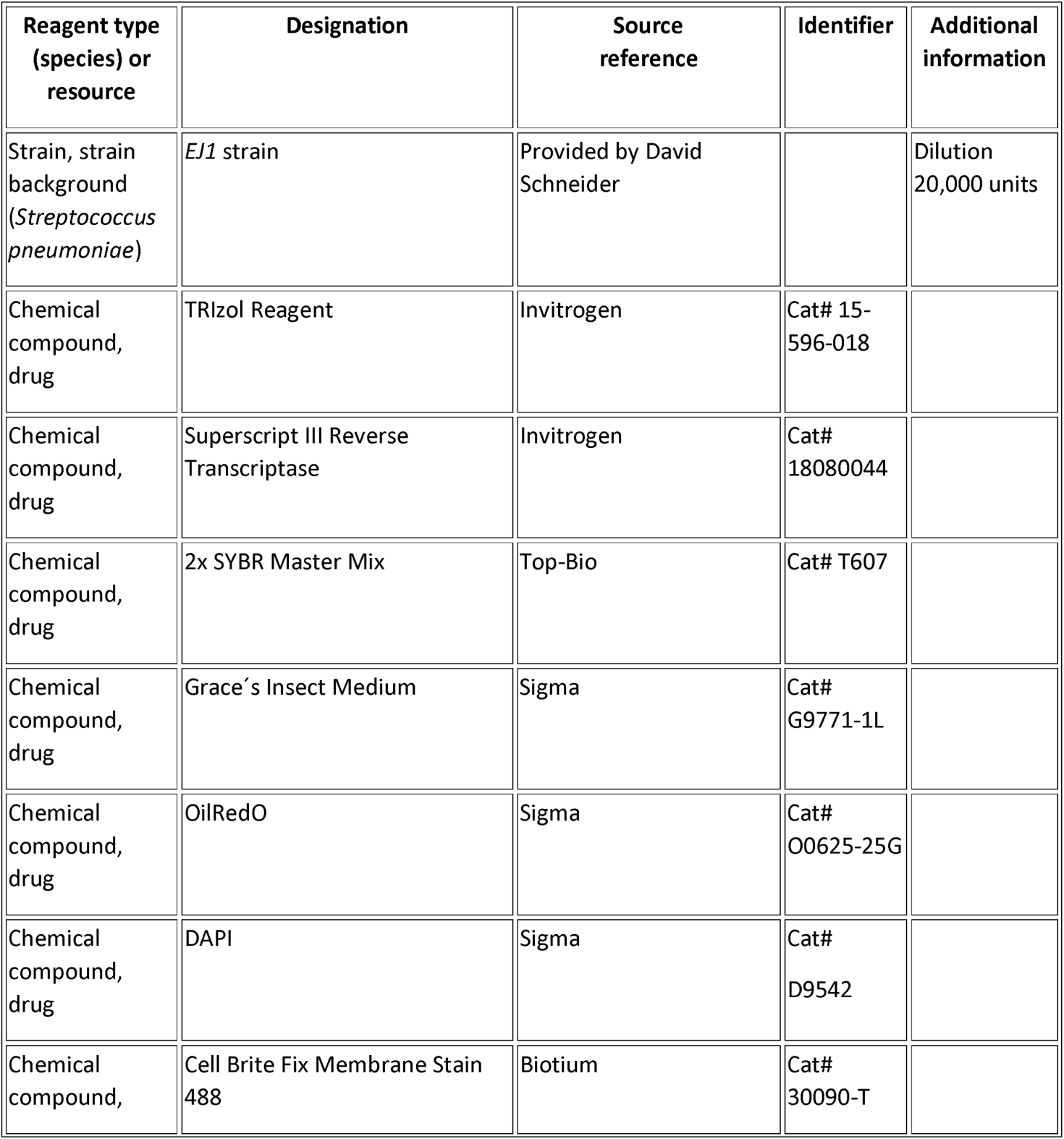
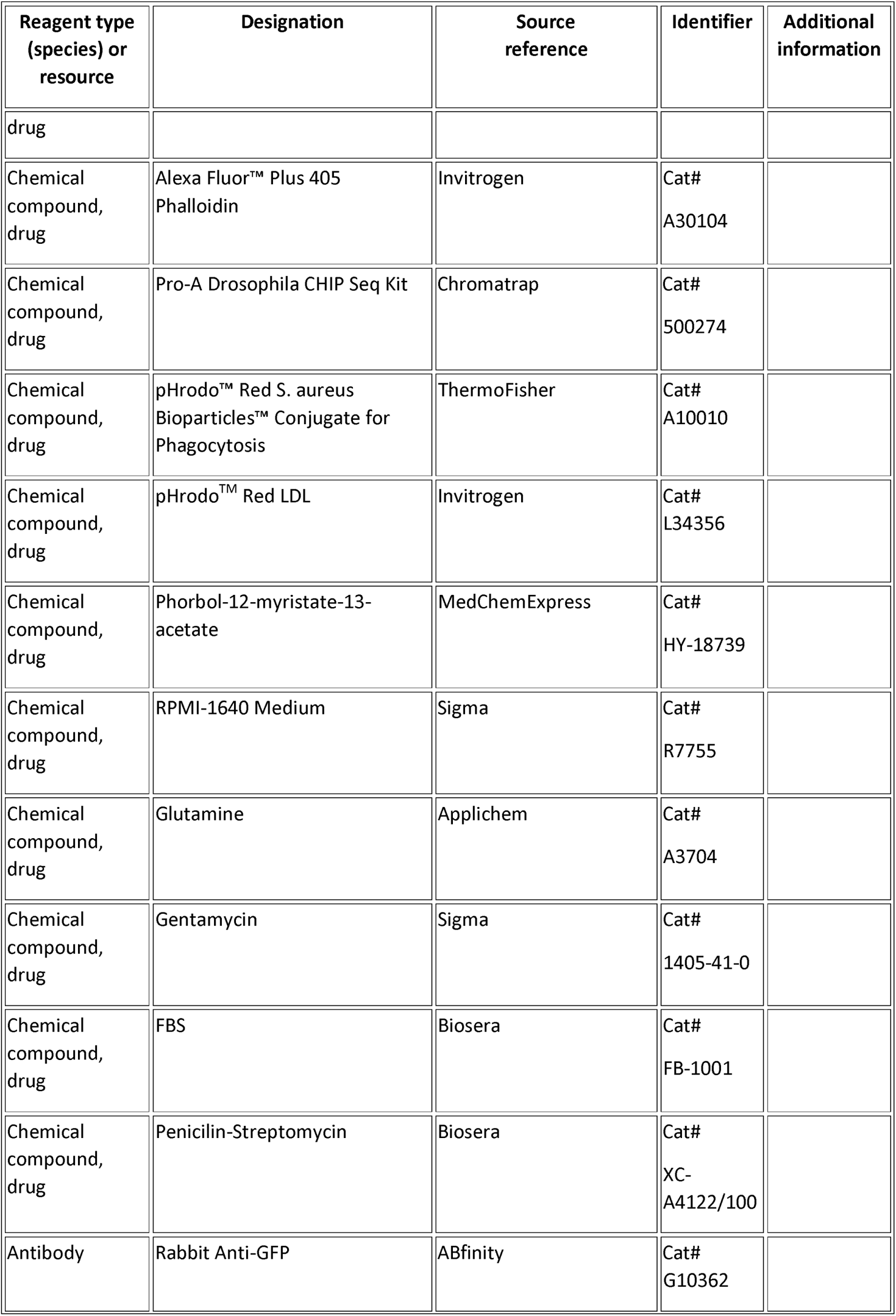
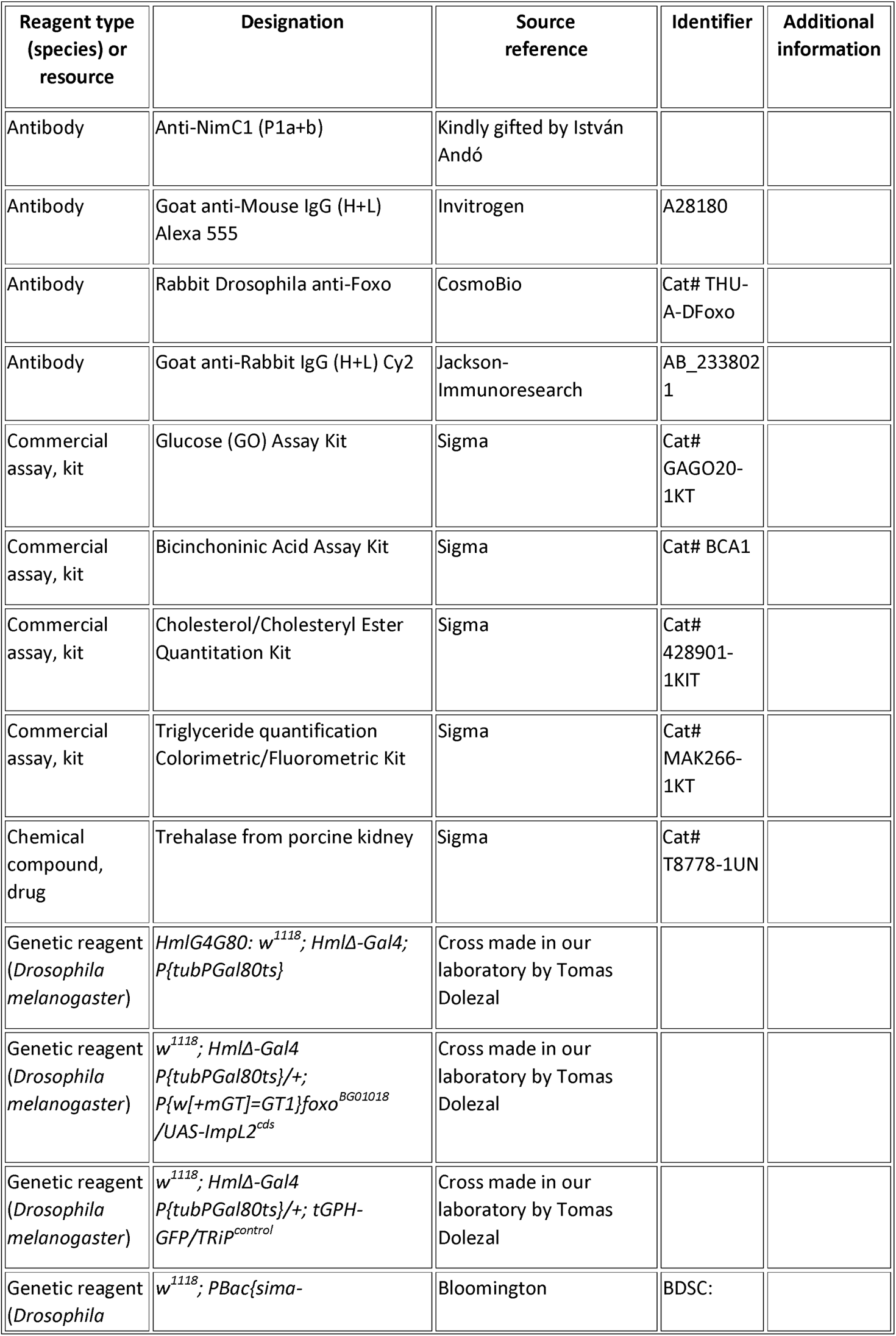
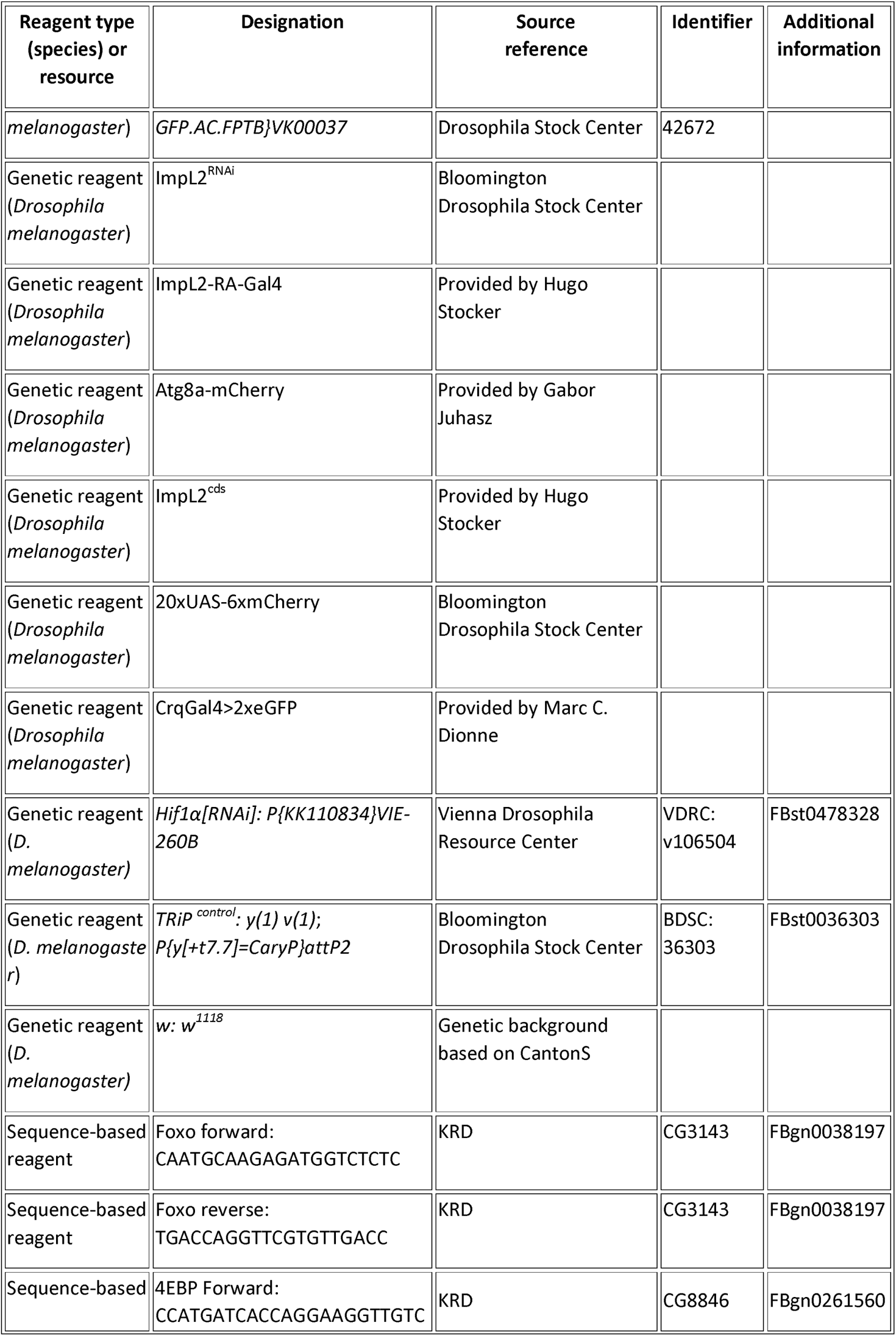
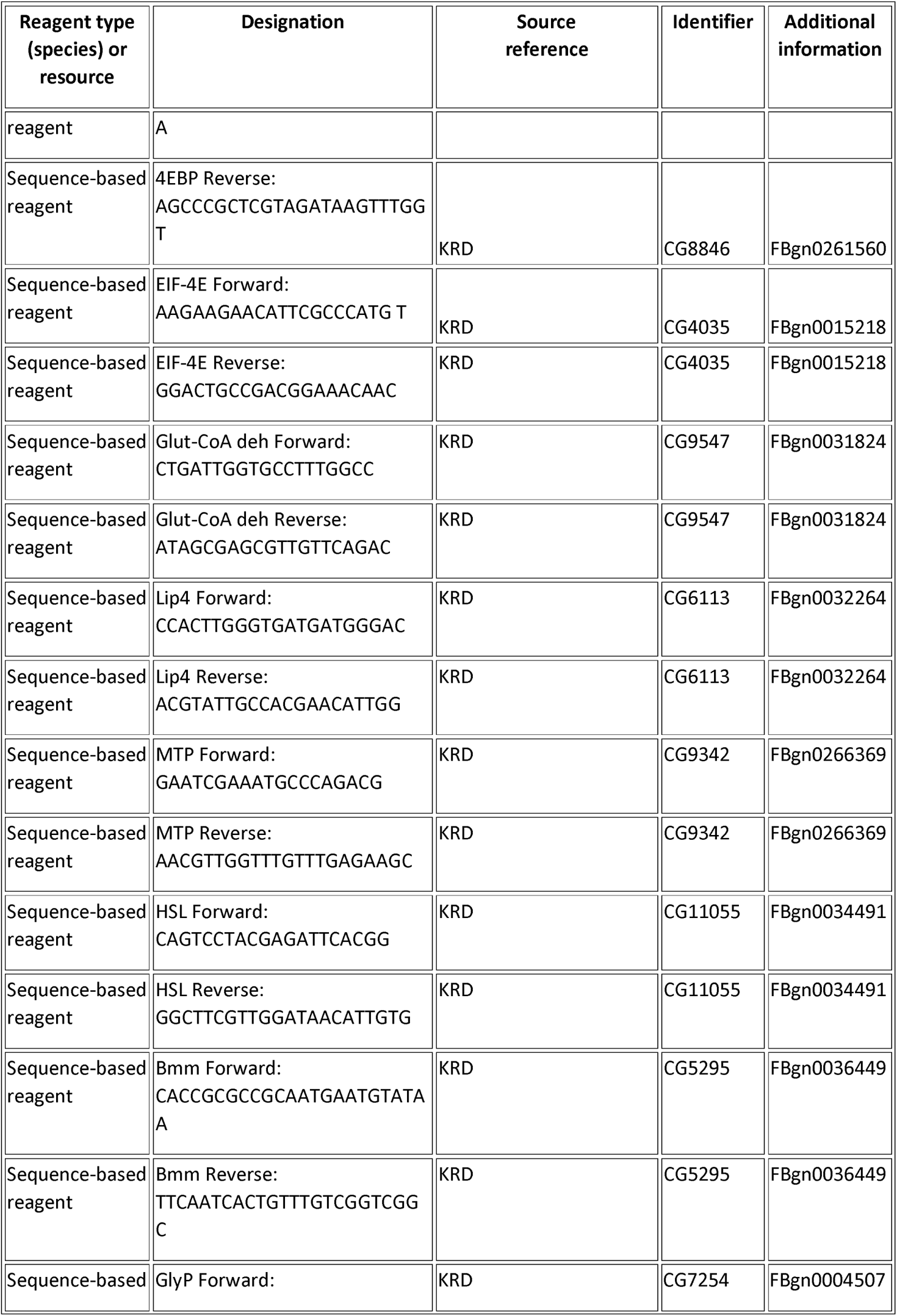
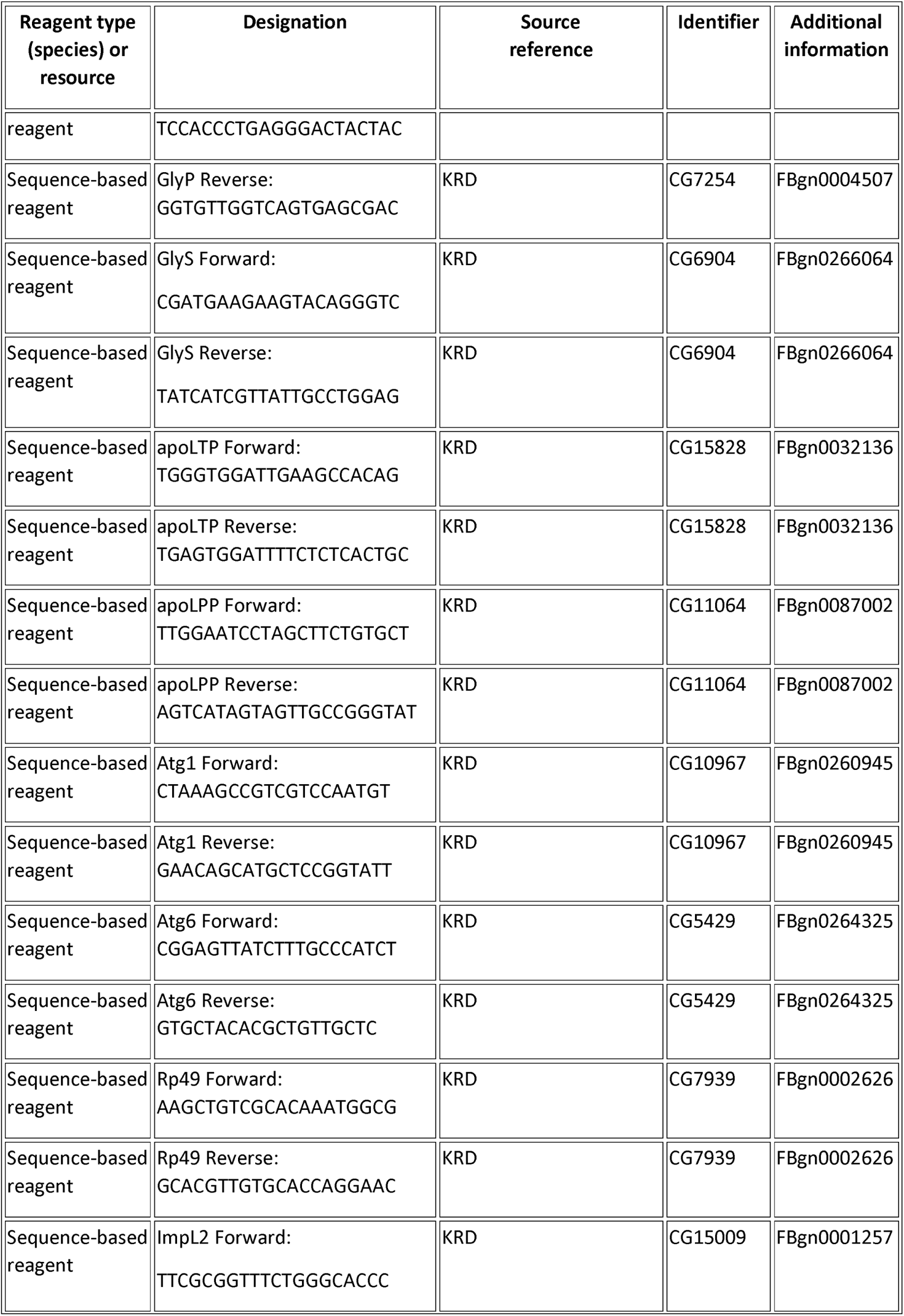
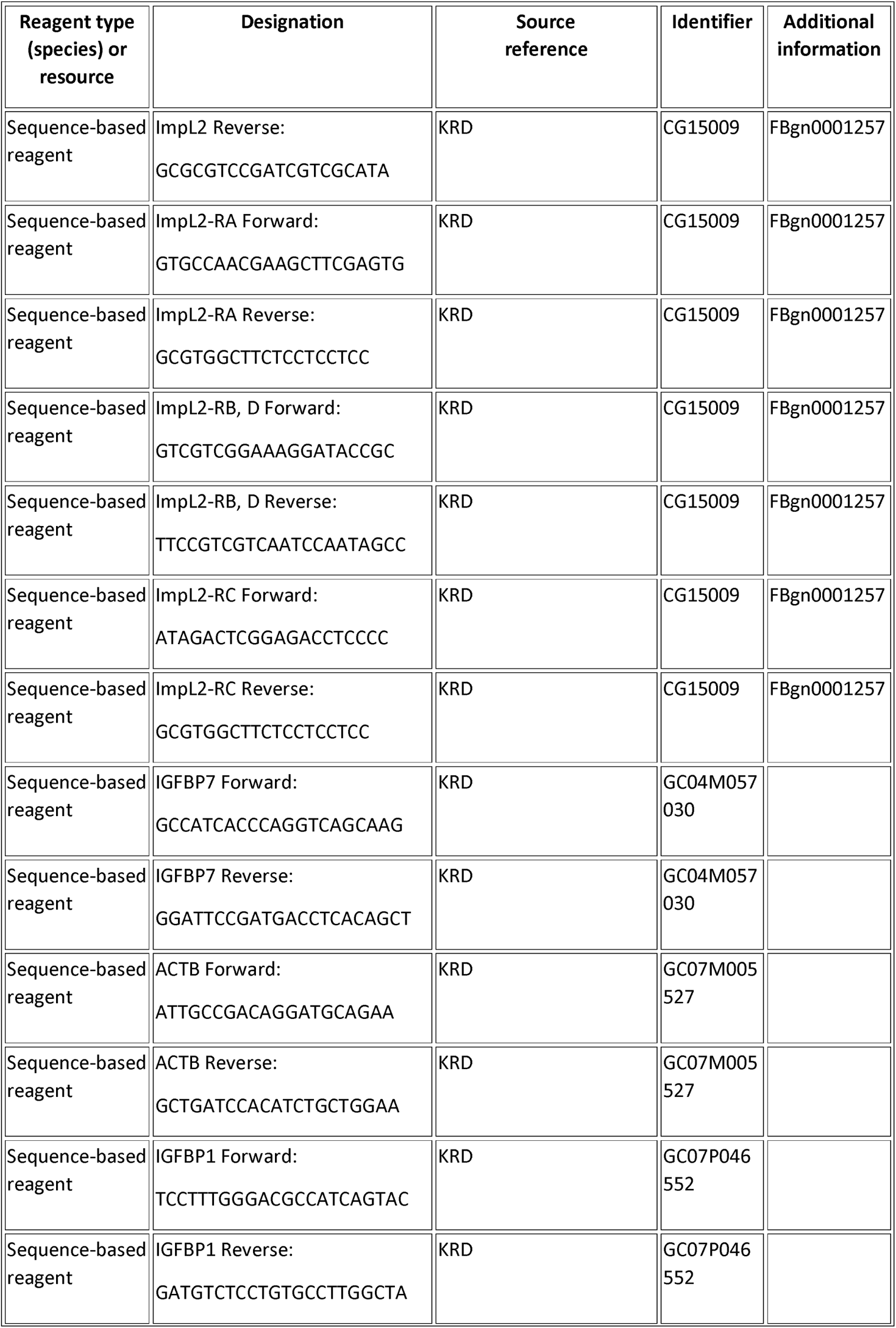
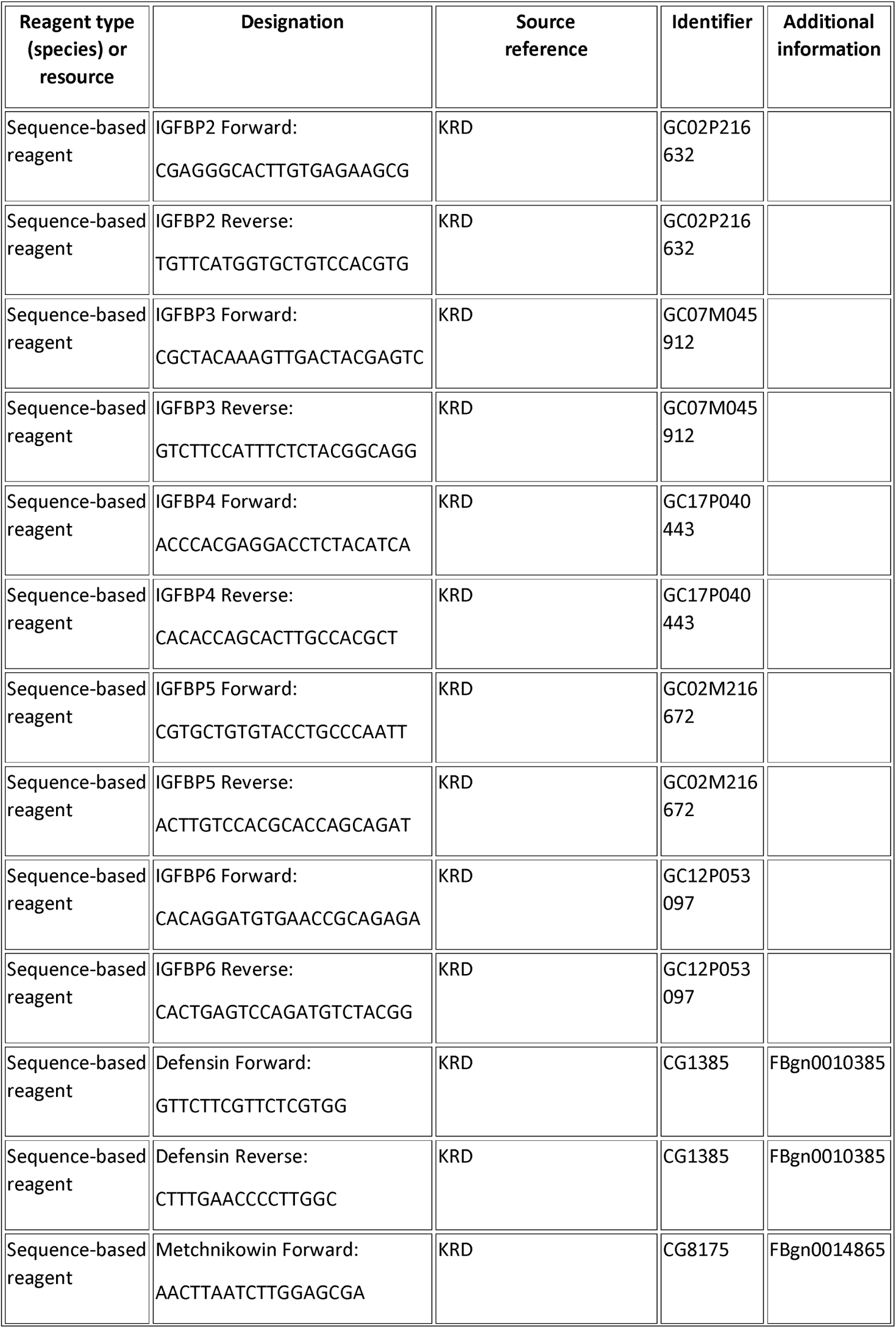
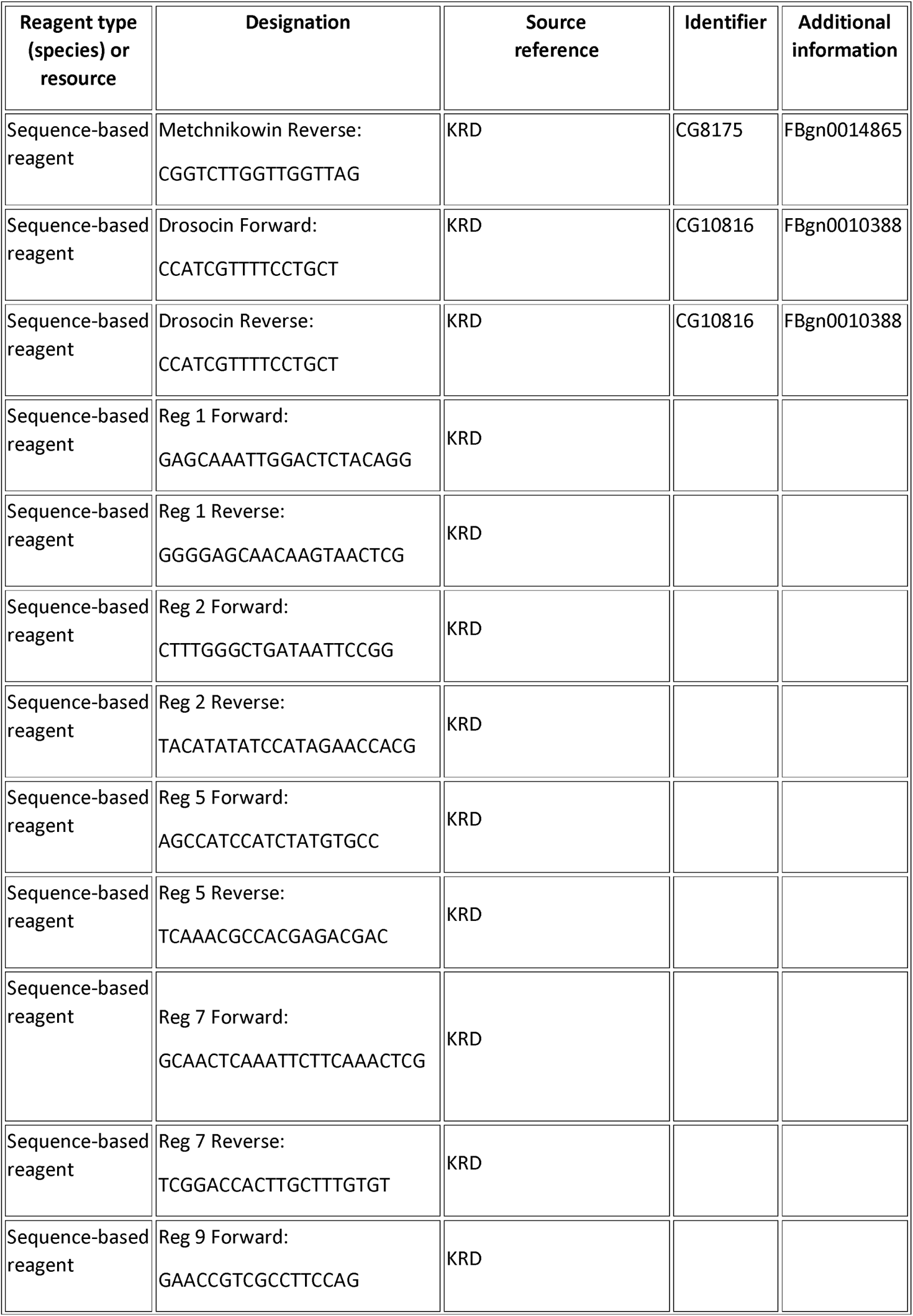
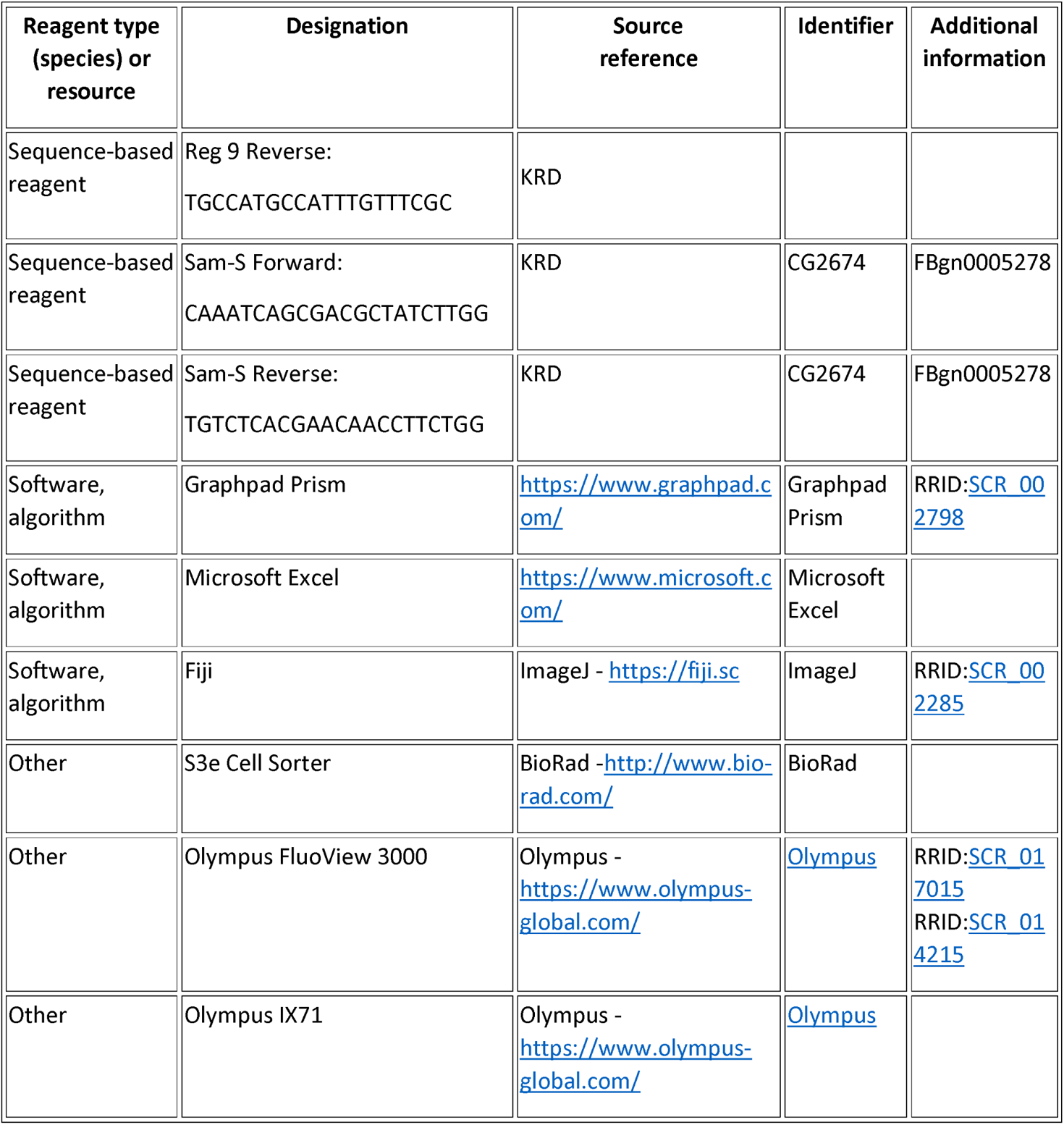

## Figure legends

Figure 1-figure supplement 1. **ImpL2-RA>mCherry-positive cells display hemocyte morphology and actively phagocytose Staphylococcus aureus**

Repesentative confocal microscopy image of ImpL2-RA>mCherry hemocyte depicting its phagocytic ability visualized by S. aureus-pHrodo (green). The scale bar represents 10 µm.

Figure 1-figure supplement 2. **Gene expression of ImpL2 transcipt variants in distinct tissues**

Gene expression of ImpL2-RA (left), ImpL2-RB and ImpL2-RD (middle) and ImpL2-RC (right) transcript variants in either hemocytes, fat body or thoracic muscles dissected from control and infected Crq>GFP flies at 24 hpi. Expression levels normalized against rp49 are presented as a fold change relative to levels of ImpL2-RA in hemocytes of Buff. injected control flies, which were arbitrarily set to 1. Individual dots represent biological replicates. Results were compared by unpaired t test with Holm-Sidak method for multiple comparisons. Values are mean ± SD, asterisks mark statistically significant differences (*p<0.05; **p<0.01; ***p<0.001).

Figure 1-figure supplement 3. **Efficiency of genetic manipulation of Hif1α**

Gene expression of Hif1α in hemocytes of Buff. injected and S.p. infected flies with or without hemocyte-specific Hif1α knockdown. Expression levels normalized against rp49 are presented as a fold change relative to levels of Hif1α in Buff. injected HmlGal>GFP flies, which were arbitrarily set to 1. Individual dots represent biological replicates. Results compared by 2way ANOVA Tukey’s multiple comparisons test. Values are mean ± SD, asterisks mark statistically significant differences (*p<0.05; **p<0.01; ***p<0.001).

Figure 1-figure supplement 4. **ImpL2-RA regulatory sequence with transcription factor binding sites**

500-bp long sequence upstream of ImpL2-RA transcriptional start site (as shown on map) with marked binding sites for Hif1α, Rel and HSF1. Hif1α, hypoxia-inducible factor 1 α; Rel, relish; HSF1, heat shock factor 1.

Figure 1-figure supplement 5. **Copy of Figure 1B and 1D in color blind friendly pallete.**

(**A, B**) Representative confocal microscopy images of control (left) and infected (right) ImpL2-RA>mCherry individuals imaged at 24 hpi, from Z stack of 7 layers, autofluorescence in the green channel was used to visualize the fly’s body. (**C**) Confocal microscopy image of infected and dissected ImpL2-RA>mCherry bearing fly stained with anti-NimC1 antibody (green) depicting the expression of ImpL2 in hemocytes, from Z stack of 11 layers. The scale bar represents 10 µm.

Figure 3-figure supplement 1. **Expression levels of genes regulating metabolism in adipose tissue**

Gene expression of Foxo, 4EBP, EIF4E1, Glutaryl-CoA deh., Lip4, HSL, Bmm, GlyS, GlyP, apoLTP, apoLPP, MTP, Atg1, and Atg6 genes in adipose tissue of Buff. injected (black bars) and S.p. infected (grey bars) flies with macrophage-specific ImpL2 knockdown, overexpression and their respective controls; all genotypes depicted below the x-axis were crossed with Hml>Gal4; TubGal80TS. Expression levels normalized against rp49 are presented as a fold-change relative to the levels in the Buff. injected TRiP^control^, which was arbitrarily set to 1. The individual dots represent biological replicates. Results were compared by 2way ANOVA Tukey’s multiple comparisons test. Bars show mean ± SD, asterisks mark statistically significant differences (*p<0.05; **p<0.01; ***p<0.001).

Figure 6-figure supplement 1. **Concentration of carbohydrates and lipids in control fly lines**

(**A-E**) Concentrations of circulating glucose (**A**), trehalose (**B**) and glycerides (**C**) in hemolymph and triglycerides (**D**) and glycogen (**E**) at a whole body level of Buff. injected and S.p. infected Hml-Gal4 TubGal80TS x TRiP^control^, Hml-Gal4 TubGal80TS x w1118, ImpL2^RNAi^ x TRiP^control^, and ImpL2^CDS^ x w1118 control flies at 24 hpi. Metabolite concentrations were normalized to the amount of proteins in each sample. Results were compared by 2way ANOVA Tukey’s multiple comparisons test. Bars show means ± SD, asterisks mark statistically significant differences (*p<0.05; **p<0.01; ***p<0.001).

Figure 7-figure supplement 1. **Survival of additional control genotypes of S. pneumoniae infection.**

Survival rate of S.p. infected and Buff. injected flies with macrophage-specific ImpL2 knockdown (ImpL2^RNAi^) (upper graph) and overexpression (ImpL2^CDS^) (lower graph) and and their respective controls (ImpL2^RNAi^ x TRiP^control^, HmlG4G80 x TRiP^control^, ImpL2cds x w1118 and HmlG4G80 x w1118). Three independent experiments were performed and combined into one survival curve; the number of individuals per replicate was at least 500 for each genotype. Survival data were analyzed by Log-rank and Grehan-Breslow Wilcoxon tests.

Figure 7-figure supplement 2. **Expression of antimicrobial peptides in hemocytes**

Gene expression of Drosocin (upper graph), Metchnikowin (middle graph) and Defensin (lower graph) in macrophages of flies with macrophage-specific ImpL2 knockdown (ImpL2^RNAi^), overexpression (ImpL2^CDS^), and their respective controls (TRiP^control^, w1118) at 24 hpi. Expression levels normalized against rp49 are reported as fold change relative to levels of Drosocin, Metchnikowin, and Defensin, respectively, in Buff. injected TRiP^control^, which were arbitrarily set to 1.The individual dots represent biological replicates. Results were compared by 2way ANOVA Tukey’s multiple comparisons test. Bars show means ± SD, asterisks mark statistically significant differences (*p<0.05; **p<0.01; ***p<0.001).

## Source Data Files

All source data files are in Graphpad Prism format, which was used for all data processing. The free Graphpad viewer is available at www.graphpad.com

Fig1.pzfx – Source data for Figure 1., Figure 1.-figure supplement 2. and Figure 1.-figure supplement 3.

Fig2.pzfx – Source data for Figure 2.

Fig3.pzfx – Source data for Figure 3. and Figure 3.-figure supplement 1. Fig6.pzfx – Source data for Figure 6.

Fig6suppl1.pzfx – Source data for Figure 6.-figure supplement 1. Fig7 – Source Data 1.pzfx – Source data for Figure 7. Fig7suppl1.pzfx – Source data for Figure 7.-figure supplement 1.

Fig6suppl2.pzfx – Source data for Figure 7.-figure supplement 2.

Fig8.pzfx – Source data for Figure 8. Fig9.pzfx – Source data for Figure 9.

## Notes

### Competing Interest Statement

The authors have declared no competing interest.

### Summary of Updates

1. Binding of Hif1α to the regulatory sequence of the ImpL2-RA transcriptional variant is demonstrated (Fig 1H). 2. Macrophage-derived ImpL2 suppresses insulin signaling in the fat body is demonstrated by tGPH reporter and rescue of ImpL2-induced effects by the foxo mutation demonstrates that the observed effects are mediated by Foxo (Fig 5). 3. Injecting flies with fluorescently labeled lipoproteins (LDL-pHRodo - Fig. 6E,F) demonstrates their uptake by macrophages. 4. All graphs are presented with individual values.

## References

Alee. 2011. “ImpL2 Represses Insulin Signaling in Response to Hypoxia.” Dissertation Thesis. University of Oregon, Eugene, Oregon, USA.

Akiel, Maaged et al. 2017. “IGFBP7 Deletion Promotes Hepatocellular Carcinoma.” Cancer Research 77(15): 4014–25. http://cancerres.aacrjournals.org/lookup/doi/10.1158/0008-5472.CAN-16-2885.

Alic, Nazif, Matthew P. Hoddinott, Giovanna Vinti, and Linda Partridge. 2011. “Lifespan Extension by Increased Expression of the Drosophila Homologue of the IGFBP7 Tumour Suppressor.” Aging Cell 10(1): 137–47. http://doi.wiley.com/10.1111/j.1474-9726.2010.00653.x.

Andrejeva, Gabriela, and Jeffrey C. Rathmell. 2017. “Similarities and Distinctions of Cancer and Immune Metabolism in Inflammation and Tumors.” Cell Metabolism 26(1): 49–70. https://linkinghub.elsevier.com/retrieve/pii/S1550413117303467.

Arquier, Nathalie et al. 2006. “Analysis of the Hypoxia-Sensing Pathway in Drosophila Melanogaster.” Biochemical Journal 393(2): 471–80. https://portlandpress.com/biochemj/article/393/2/471/78893/Analysis-of-the-hypoxiasensing-pathway-in.

Aspichueta, Patricia et al. 2012. “Disrupted VLDL Features and Lipoprotein Metabolism in Sepsis.” In *Dyslipidemia - From Prevention to Treatment*, InTech. http://www.intechopen.com/books/dyslipidemia-from-prevention-to-treatment/disrupted-vldl-features-and-lipoprotein-metabolism-in-sepsis.

Bader, R. et al. 2013. “The IGFBP7 Homolog Imp-L2 Promotes Insulin Signaling in Distinct Neurons of the Drosophila Brain.” Journal of Cell Science 126(12): 2571–76. http://jcs.biologists.org/cgi/doi/10.1242/jcs.120261.

Bajgar, Adam et al. 2015. “Extracellular Adenosine Mediates a Systemic Metabolic Switch during Immune Response” ed. Marc S. Dionne. PLOS Biology 13(4): e1002135. https://dx.plos.org/10.1371/journal.pbio.1002135.

Bajgar, Adam, and Tomas Dolezal. 2018. “Extracellular Adenosine Modulates Host-Pathogen Interactions through Regulation of Systemic Metabolism during Immune Response in Drosophila” ed. Brian P. Lazzaro. PLOS Pathogens 14(4): e1007022. https://dx.plos.org/10.1371/journal.ppat.1007022.

Bayley, Jeppe Seamus et al. 2020. “Cold-Acclimation Increases Depolarization Resistance and Tolerance in Muscle Fibers from a Chill-Susceptible Insect, Locusta Migratoria.” *American Journal of Physiology-Regulatory*, Integrative and Comparative Physiology: ajpregu.00068.2020. https://journals.physiology.org/doi/10.1152/ajpregu.00068.2020.

Biswas, Subhra K., and Alberto Mantovani. 2012. “Orchestration of Metabolism by Macrophages.” Cell Metabolism 15(4): 432–37. https://linkinghub.elsevier.com/retrieve/pii/S1550413112000150.

Van den Bossche, Jan, Luke A. O’Neill, and Deepthi Menon. 2017. “Macrophage Immunometabolism: Where Are We (Going)?” Trends in Immunology 38(6): 395–406. https://linkinghub.elsevier.com/retrieve/pii/S147149061730042X.

Bunker, Brandon D et al. 2015. “The Transcriptional Response to Tumorigenic Polarity Loss in Drosophila.” eLife 4. http://www.ncbi.nlm.nih.gov/pubmed/25719210.

Chen, Grischa Y., Daniel A. Pensinger, and John-Demian Sauer. 2017. “Listeria Monocytogenes Cytosolic Metabolism Promotes Replication, Survival, and Evasion of Innate Immunity.” Cellular Microbiology 19(10): e12762. https://onlinelibrary.wiley.com/doi/10.1111/cmi.12762.

Chistiakov, Dimitry A. et al. 2017. “Mechanisms of Foam Cell Formation in Atherosclerosis.” Journal of Molecular Medicine 95(11): 1153–65. http://link.springer.com/10.1007/s00109-017-1575-8.

Clark, Rebecca I. et al. 2011. “Multiple TGF-β Superfamily Signals Modulate the Adult Drosophila Immune Response.” Current Biology 21(19): 1672–77. https://linkinghub.elsevier.com/retrieve/pii/S0960982211009547.

Dev, R, E. Bruera, and S. Dalal. 2018. “Insulin Resistance and Body Composition in Cancer Patients.” Annals of Oncology 29: ii18–26. https://linkinghub.elsevier.com/retrieve/pii/S0923753419316801.

Dionne, Marc S., Linh N. Pham, Mimi Shirasu-Hiza, and David S. Schneider. 2006. “Akt and Foxo Dysregulation Contribute to Infection-Induced Wasting in Drosophila.” Current Biology 16(20): 1977–85. https://linkinghub.elsevier.com/retrieve/pii/S0960982206020641.

Dolezal, Tomas et al. 2019. “Molecular Regulations of Metabolism during Immune Response in Insects.” Insect Biochemistry and Molecular Biology 109: 31–42. https://linkinghub.elsevier.com/retrieve/pii/S0965174818304600.

Febbraio, Maria, Ella Guy, and Roy L. Silverstein. 2004. “Stem Cell Transplantation Reveals That Absence of Macrophage CD36 Is Protective Against Atherosclerosis.” *Arteriosclerosis*, Thrombosis, and Vascular Biology 24(12): 2333–38. https://www.ahajournals.org/doi/10.1161/01.ATV.0000148007.06370.68.

Figueroa-Clarevega, Alejandra, and David Bilder. 2015. “Malignant Drosophila Tumors Interrupt Insulin Signaling to Induce Cachexia-like Wasting.” Developmental Cell 33(1): 47–55. https://linkinghub.elsevier.com/retrieve/pii/S1534580715001434.

Gilbert, Lawrence I., and Haruo Chino. 1974. “Transport of Lipids in Insects.” Journal of Lipid Research 15(5): 439–56. https://linkinghub.elsevier.com/retrieve/pii/S002222752036764X.

Gunnerson, Kyle J. et al. 2016. “TIMP2•IGFBP7 Biomarker Panel Accurately Predicts Acute Kidney Injury in High-Risk Surgical Patients.” Journal of Trauma and Acute Care Surgery 80(2): 243–49. http://journals.lww.com/01586154-201602000-00009.

H. Tomkin, Gerald. 2012. “LDL as a Cause of Atherosclerosis.” The Open Atherosclerosis & Thrombosis Journal 5(1): 13–21. http://benthamopen.com/ABSTRACT/TOATHERTJ-5-13.

Harris, H W, J E Gosnell, and Z L Kumwenda. 2000. “The Lipemia of Sepsis: Triglyceride-Rich Lipoproteins as Agents of Innate Immunity.” Journal of endotoxin research 6(6): 421–30. http://www.ncbi.nlm.nih.gov/pubmed/11521066.

Honegger, Basil et al. 2008. “Imp-L2, a Putative Homolog of Vertebrate IGF-Binding Protein 7, Counteracts Insulin Signaling in Drosophila and Is Essential for Starvation Resistance.” Journal of Biology 7(3): 10. http://jbiol.biomedcentral.com/articles/10.1186/jbiol72.

Kamps-Hughes, Nick, Jessica L. Preston, Melissa A. Randel, and Eric A. Johnson. 2015. “Genome-Wide Identification of Hypoxia-Induced Enhancer Regions.” PeerJ 3: e1527. https://peerj.com/articles/1527.

Kelly, Beth, and Luke AJ O’Neill. 2015. “Metabolic Reprogramming in Macrophages and Dendritic Cells in Innate Immunity.” Cell Research 25(7): 771–84. http://www.nature.com/articles/cr201568.

Khovidhunkit, Weerapan et al. 2004. “Thematic Review Series: The Pathogenesis of Atherosclerosis. Effects of Infection and Inflammation on Lipid and Lipoprotein Metabolism Mechanisms and Consequences to the Host.” Journal of Lipid Research 45(7): 1169–96. http://www.jlr.org/lookup/doi/10.1194/jlr.R300019-JLR200.

Krejčová, Gabriela et al. 2019. “Drosophila Macrophages Switch to Aerobic Glycolysis to Mount Effective Antibacterial Defense.” eLife 8. https://elifesciences.org/articles/50414.

Kwon, Young et al. 2015. “Systemic Organ Wasting Induced by Localized Expression of the Secreted Insulin/IGF Antagonist ImpL2.” Developmental Cell 33(1): 36–46. https://linkinghub.elsevier.com/retrieve/pii/S1534580715001148.

Li, Yan et al. 2013. “HIF- and Non-HIF-Regulated Hypoxic Responses Require the Estrogen-Related Receptor in Drosophila Melanogaster” ed. Eric Rulifson. PLoS Genetics 9(1): e1003230. https://dx.plos.org/10.1371/journal.pgen.1003230.

Liu, Yi et al. 2015. “Serum IGFBP7 Levels Associate with Insulin Resistance and the Risk of Metabolic Syndrome in a Chinese Population.” Scientific Reports 5(1): 10227. http://www.nature.com/articles/srep10227.

Louie, Alexander et al. 2016. “How Many Parameters Does It Take to Describe Disease Tolerance?” ed. Andy P. Dobson. PLOS Biology 14(4): e1002435. https://dx.plos.org/10.1371/journal.pbio.1002435.

Luong, Nancy et al. 2006. “Activated FOXO-Mediated Insulin Resistance Is Blocked by Reduction of TOR Activity.” Cell Metabolism 4(2): 133–42. https://linkinghub.elsevier.com/retrieve/pii/S1550413106002348.

Martínez-Castillo, Moisés et al. 2020. “Differential Production of Insulin-like Growth Factor-Binding Proteins in Liver Fibrosis Progression.” Molecular and Cellular Biochemistry 469(1–2): 65–75. http://link.springer.com/10.1007/s11010-020-03728-4.

Mohanraj L, Kim H-S, Li W, Cai Q, Kim KE, Shin H-J, et al. 2013. IGFBP-3 Inhibits Cytokine-Induced Insulin Resistance and Early Manifestations of Atherosclerosis. PLoS ONE 8(1): e55084. https://doi.org/10.1371/journal.pone.0055084

Molaei, Maral, Crissie Vandehoef, and Jason Karpac. 2019. “NF-ΚB Shapes Metabolic Adaptation by Attenuating Foxo-Mediated Lipolysis in Drosophila.” Developmental Cell 49(5): 802–810.e6. https://linkinghub.elsevier.com/retrieve/pii/S1534580719302795.

Morgantini, Cecilia et al. 2019. “Liver Macrophages Regulate Systemic Metabolism through Non-Inflammatory Factors.” Nature Metabolism 1(4): 445–59. http://www.nature.com/articles/s42255-019-0044-9.

Nagao, Ayako et al. 2019. “HIF-1-Dependent Reprogramming of Glucose Metabolic Pathway of Cancer Cells and Its Therapeutic Significance.” International Journal of Molecular Sciences 20(2): 238. https://www.mdpi.com/1422-0067/20/2/238.

Newsholme, P, R Curi, S Gordon, and E A Newsholme. 1986. “Metabolism of Glucose, Glutamine, Long-Chain Fatty Acids and Ketone Bodies by Murine Macrophages.” Biochemical Journal 239(1): 121–25. https://portlandpress.com/biochemj/article/239/1/121/21648/Metabolism-of-glucose-glutamine-longchain-fatty.

Nicholson, David, and Lindsay B. Nicholson. 2008. “A Simple Immune System Simulation Reveals Optimal Movement and Cell Density Parameters for Successful Target Clearance.” Immunology 123(4): 519–27. http://doi.wiley.com/10.1111/j.1365-2567.2007.02721.x.

Odegaard, J. I., and A. Chawla. 2013. “Pleiotropic Actions of Insulin Resistance and Inflammation in Metabolic Homeostasis.” Science 339(6116): 172–77. https://www.sciencemag.org/lookup/doi/10.1126/science.1230721.

Oh, Youngman et al. 1996. “Synthesis and Characterization of Insulin-like Growth Factor-Binding Protein (IGFBP)-7.” Journal of Biological Chemistry 271(48): 30322–25. http://www.jbc.org/lookup/doi/10.1074/jbc.271.48.30322.

Okamoto, N. et al. 2013. “A Secreted Decoy of InR Antagonizes Insulin/IGF Signaling to Restrict Body Growth in Drosophila.” Genes & Development 27(1): 87–97. http://genesdev.cshlp.org/cgi/doi/10.1101/gad.204479.112.

Owusu-Ansah, Edward, Wei Song, and Norbert Perrimon. 2013. “Muscle Mitohormesis Promotes Longevity via Systemic Repression of Insulin Signaling.” Cell 155(3): 699–712. https://linkinghub.elsevier.com/retrieve/pii/S0092867413011586.

Péan, Claire B. et al. 2017. “Regulation of Phagocyte Triglyceride by a STAT-ATG2 Pathway Controls Mycobacterial Infection.” Nature Communications 8(1): 14642. http://www.nature.com/articles/ncomms14642.

Podrez, Eugene A. et al. 2000. “Macrophage Scavenger Receptor CD36 Is the Major Receptor for LDL Modified by Monocyte-Generated Reactive Nitrogen Species.” Journal of Clinical Investigation 105(8): 1095–1108. http://www.jci.org/articles/view/8574.

Remmerie, Anneleen, and Charlotte L. Scott. 2018. “Macrophages and Lipid Metabolism.” Cellular Immunology 330: 27–42. https://linkinghub.elsevier.com/retrieve/pii/S0008874918300327.

Ruan, Wenjing et al. 2017. “Serum Levels of IGFBP7 Are Elevated during Acute Exacerbation in COPD Patients.” International Journal of Chronic Obstructive Pulmonary Disease Volume 12: 1775–80. https://www.dovepress.com/serum-levels-of-igfbp7-are-elevated-during-acute-exacerbation-in-copd--peer-reviewed-article-COPD.

Shattuck-Heidorn, Heather et al. 2016. “Energetics and the Immune System: Trade-Offs Associated with Non-Acute Levels of CRP in Adolescent Gambian Girls.” Evolution, Medicine, and Public Health: eow034. https://academic.oup.com/emph/article-lookup/doi/10.1093/emph/eow034.

Soeters, Maarten R., and Peter B. Soeters. 2012. “The Evolutionary Benefit of Insulin Resistance.” Clinical Nutrition 31(6): 1002–7. https://linkinghub.elsevier.com/retrieve/pii/S0261561412001112.

Straub, Rainer H. 2014. “Insulin Resistance, Selfish Brain, and Selfish Immune System: An Evolutionarily Positively Selected Program Used in Chronic Inflammatory Diseases.” Arthritis Research & Therapy 16(Suppl 2): S4. http://arthritis-research.biomedcentral.com/articles/10.1186/ar4688.

Teng, Ooiean, Candice Ke En Ang, and Xue Li Guan. 2017. “Macrophage–Bacteria Interactions—A Lipid-Centric Relationship.” Frontiers in Immunology 8. http://journal.frontiersin.org/article/10.3389/fimmu.2017.01836/full.

Warburg, Otto. 1925. “Über Den Stoffwechsel Der Carcinomzelle.” Klinische Wochenschrift 4(12): 534–36. http://link.springer.com/10.1007/BF01726151.

Zmora, Niv, Stavros Bashiardes, Maayan Levy, and Eran Elinav. 2017. “The Role of the Immune System in Metabolic Health and Disease.” Cell Metabolism 25(3): 506–21. https://linkinghub.elsevier.com/retrieve/pii/S1550413117300967.

