## Supplementary figures and images for "Macrophage-derived insulin/IGF antagonist ImpL2 regulates systemic metabolism for mounting an effective acute immune response in *Drosophila*"

### Figure 1-figure supplement 1

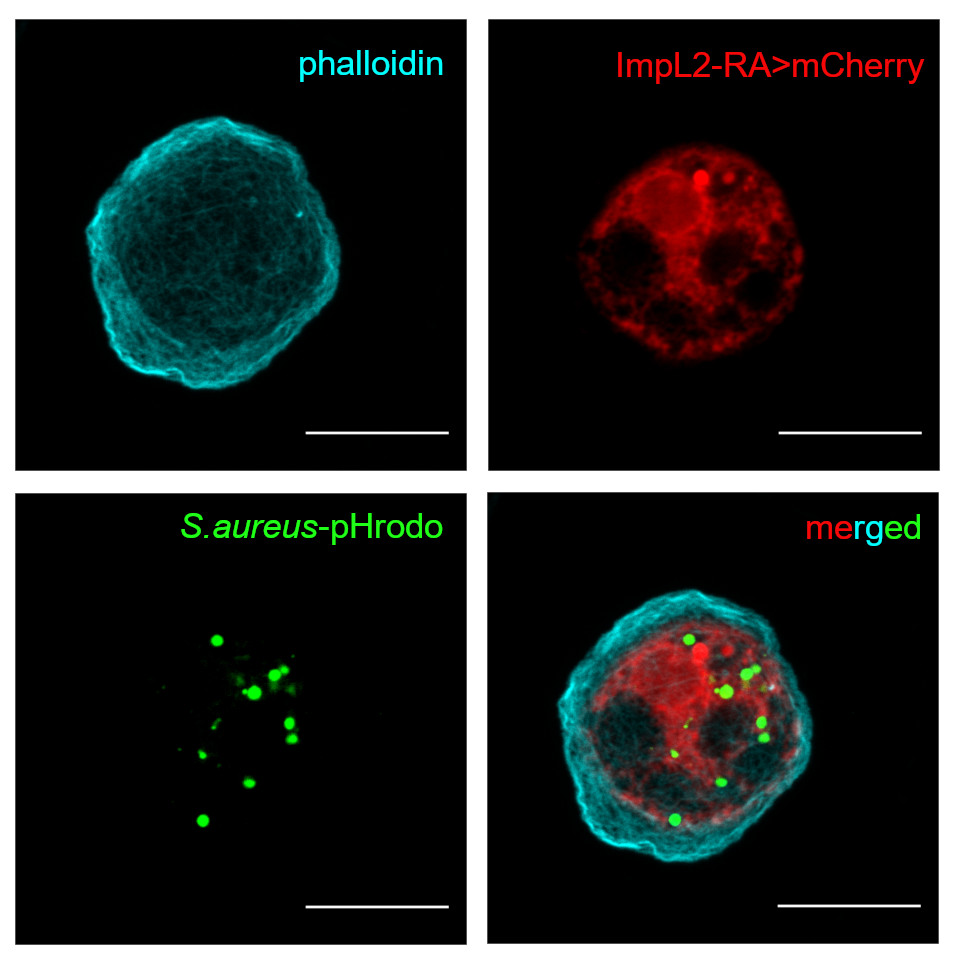

### Figure 1-figure supplement 2

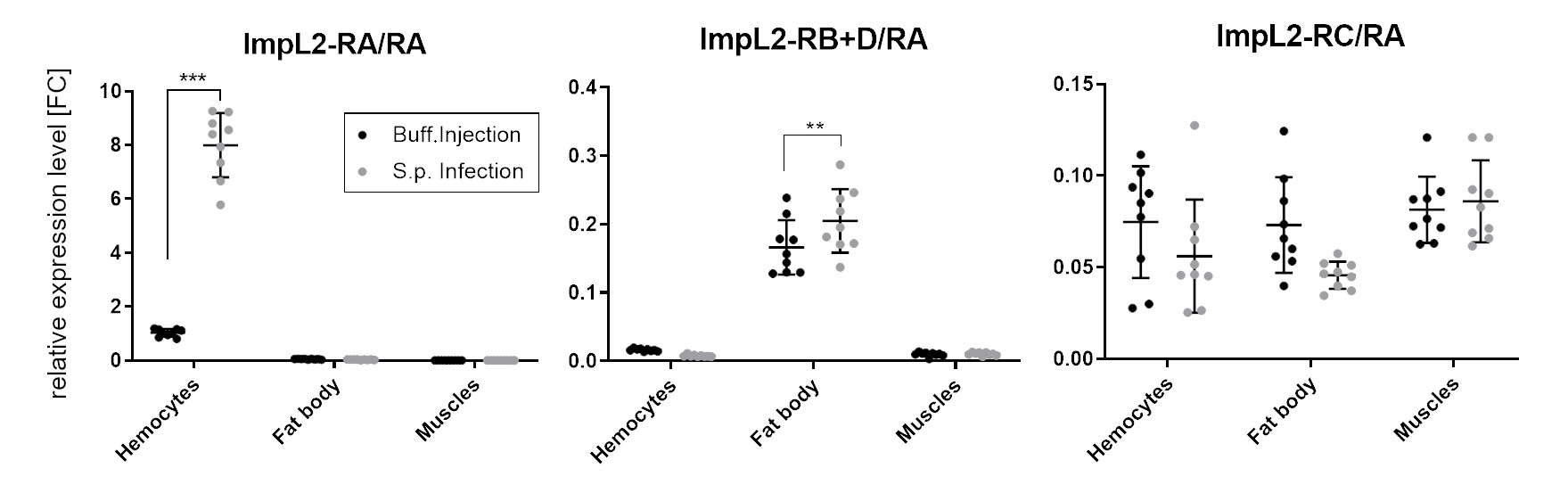

### Figure 1-figure supplement 3

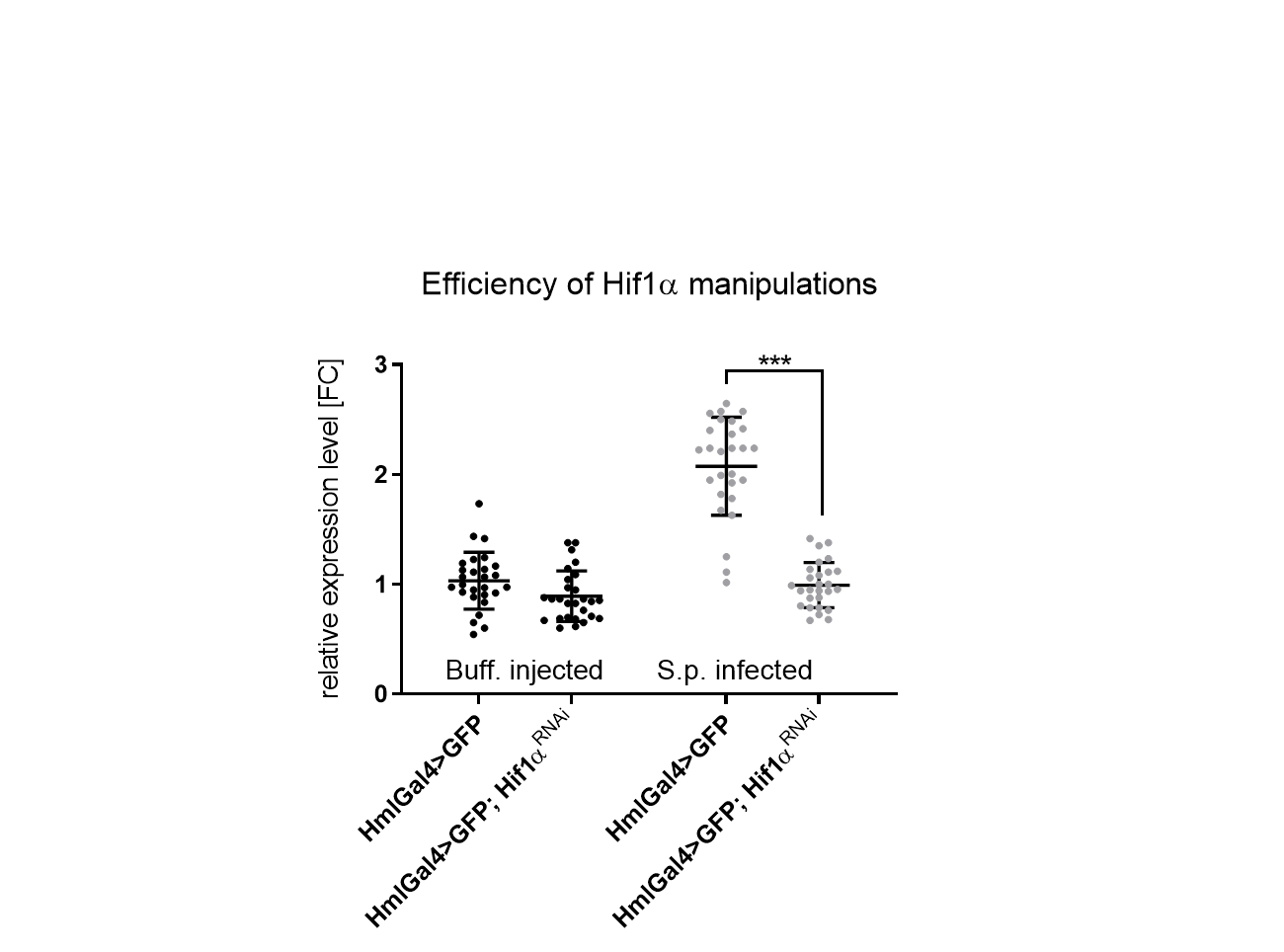

### Figure 1-figure supplement 4

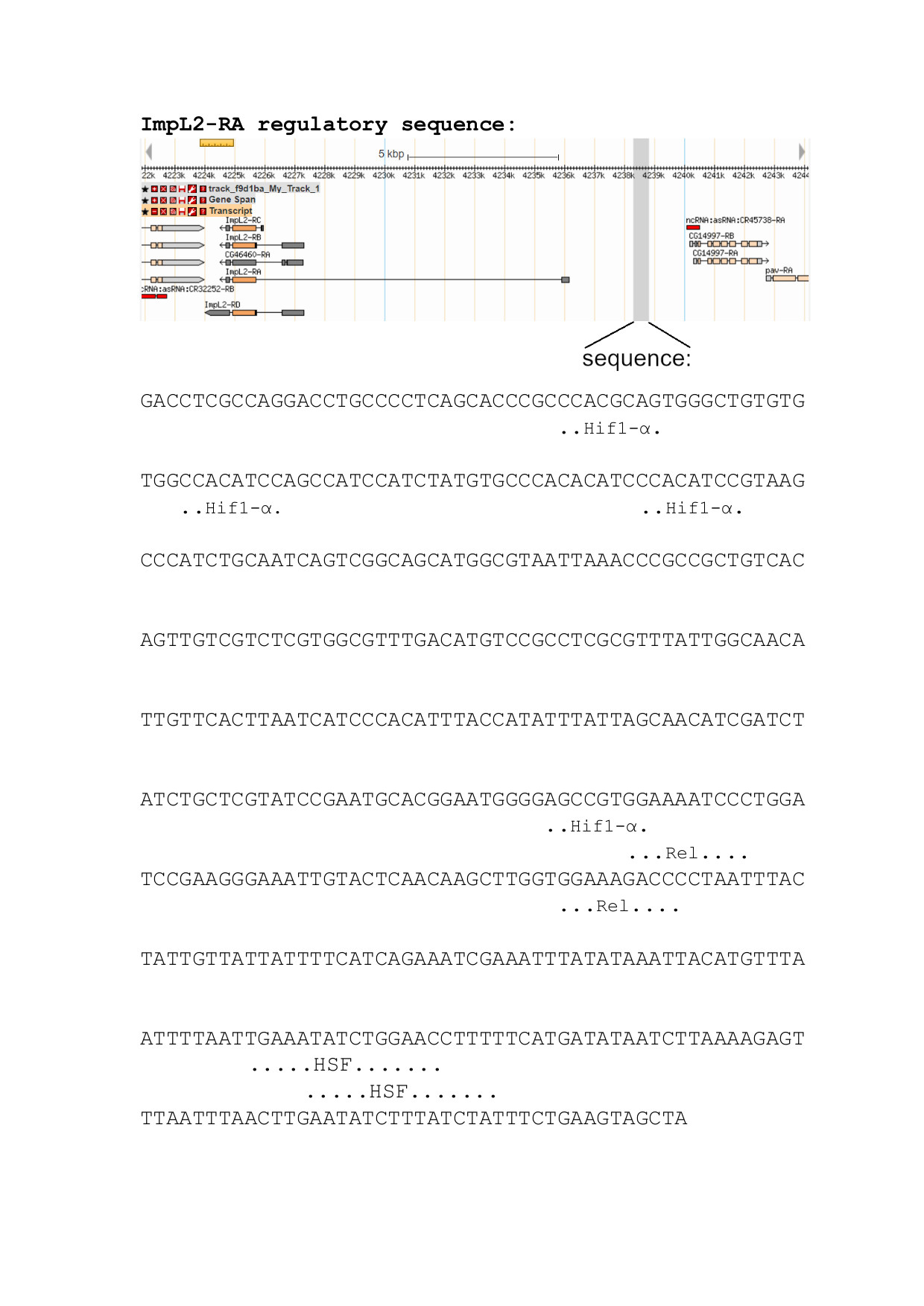

### Figure 1-figure supplement 5

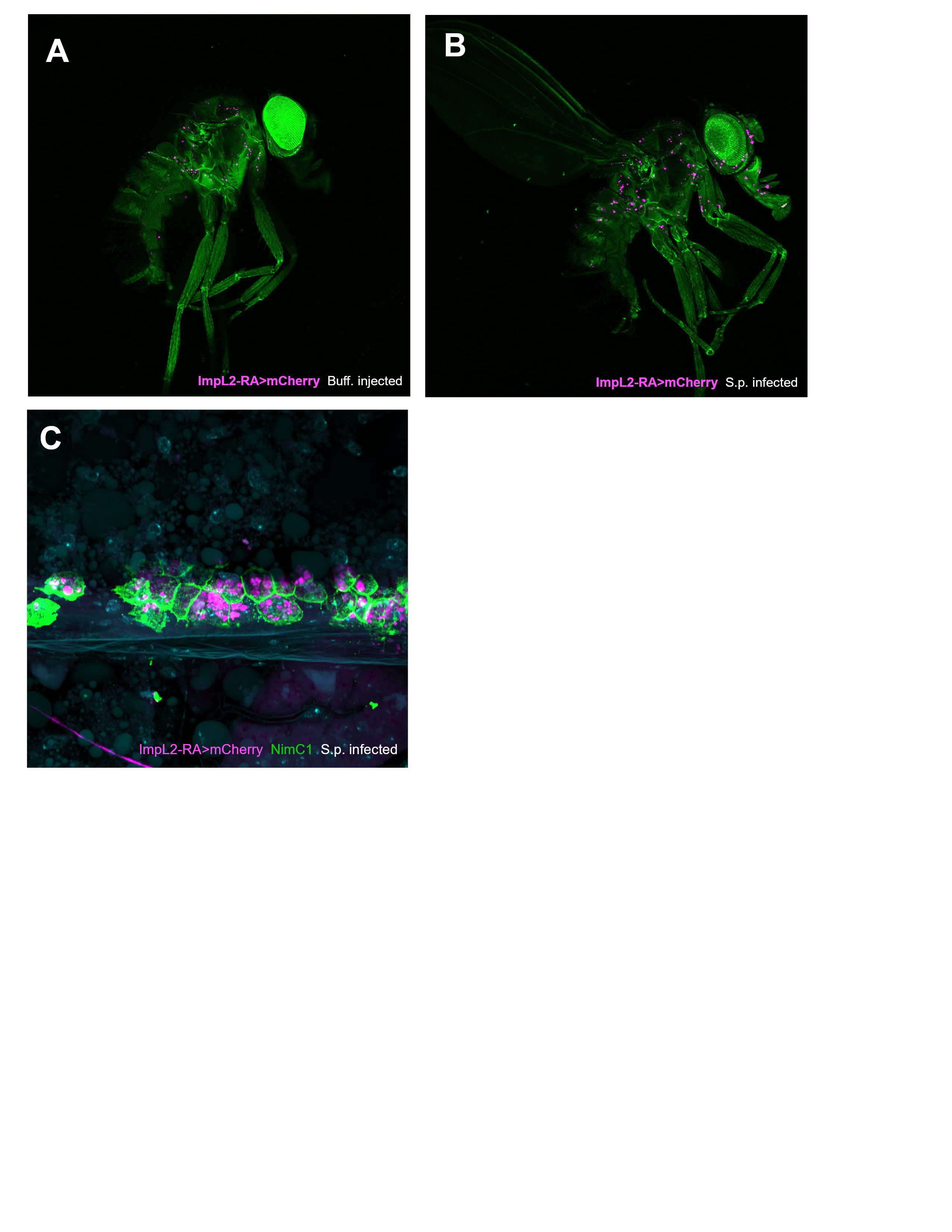

### Figure 3-figure supplement 1

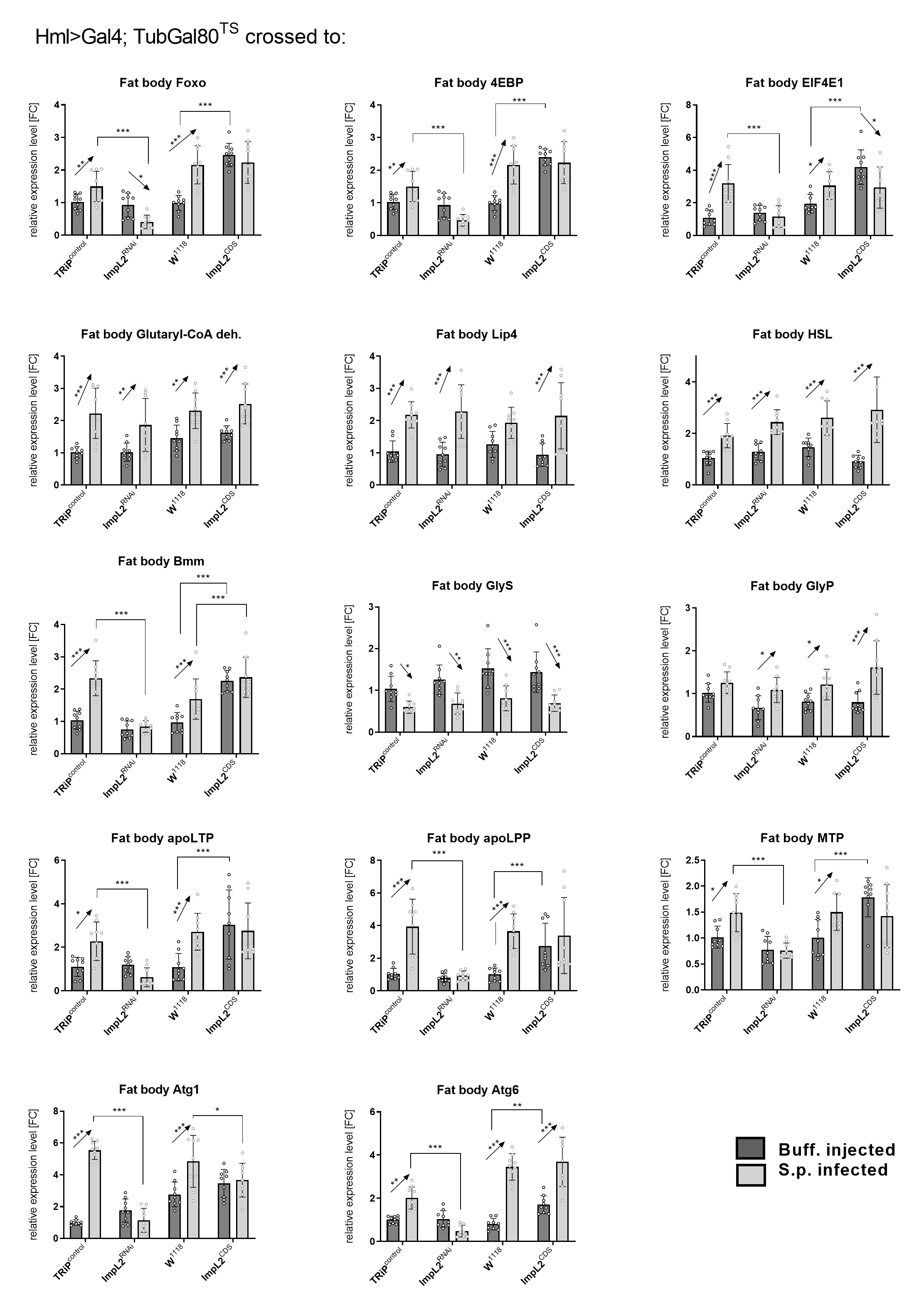

### Figure 6-figure supplement 1

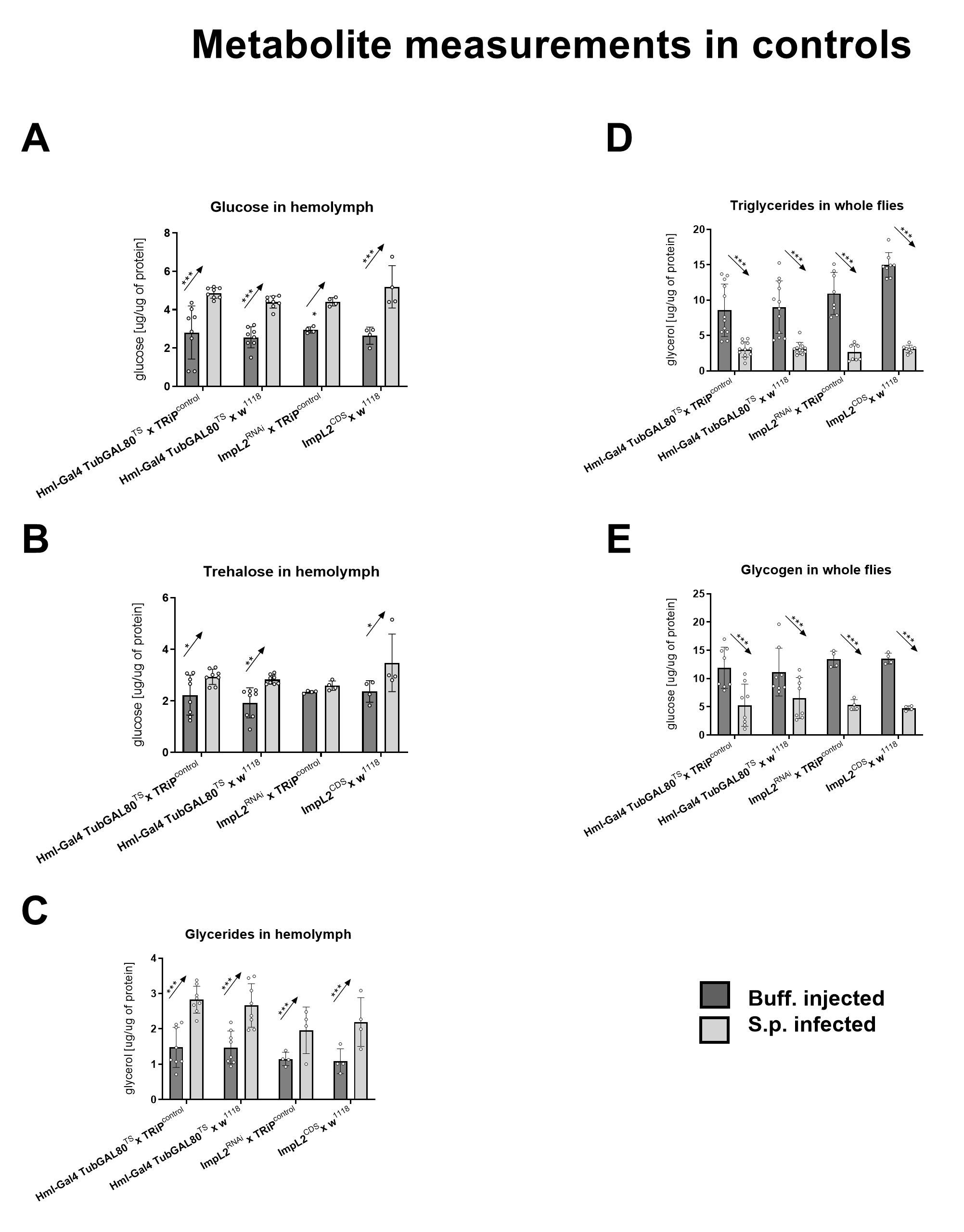

### Figure 7-figure supplement 1

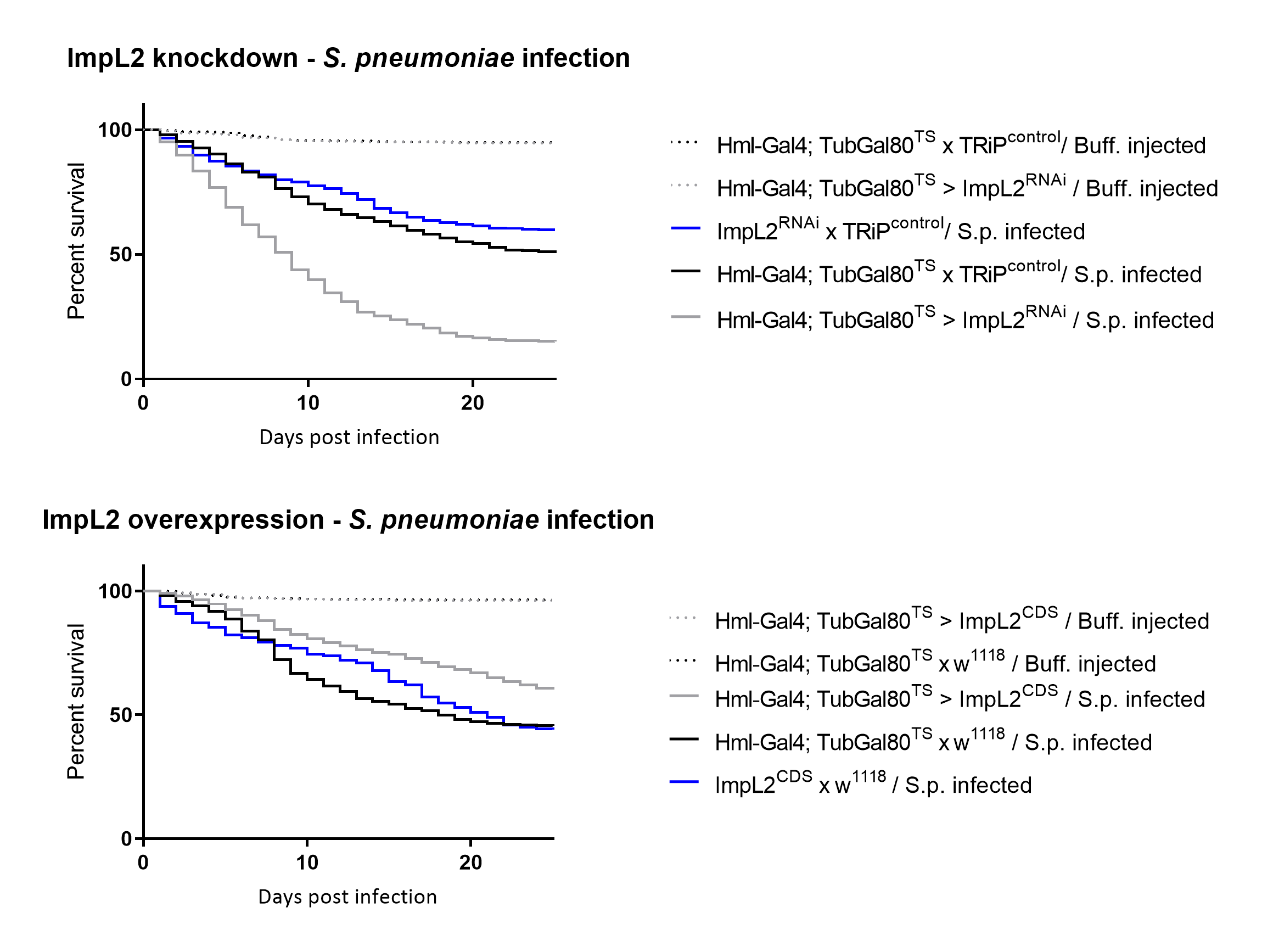

### Figure 7-figure supplement 2

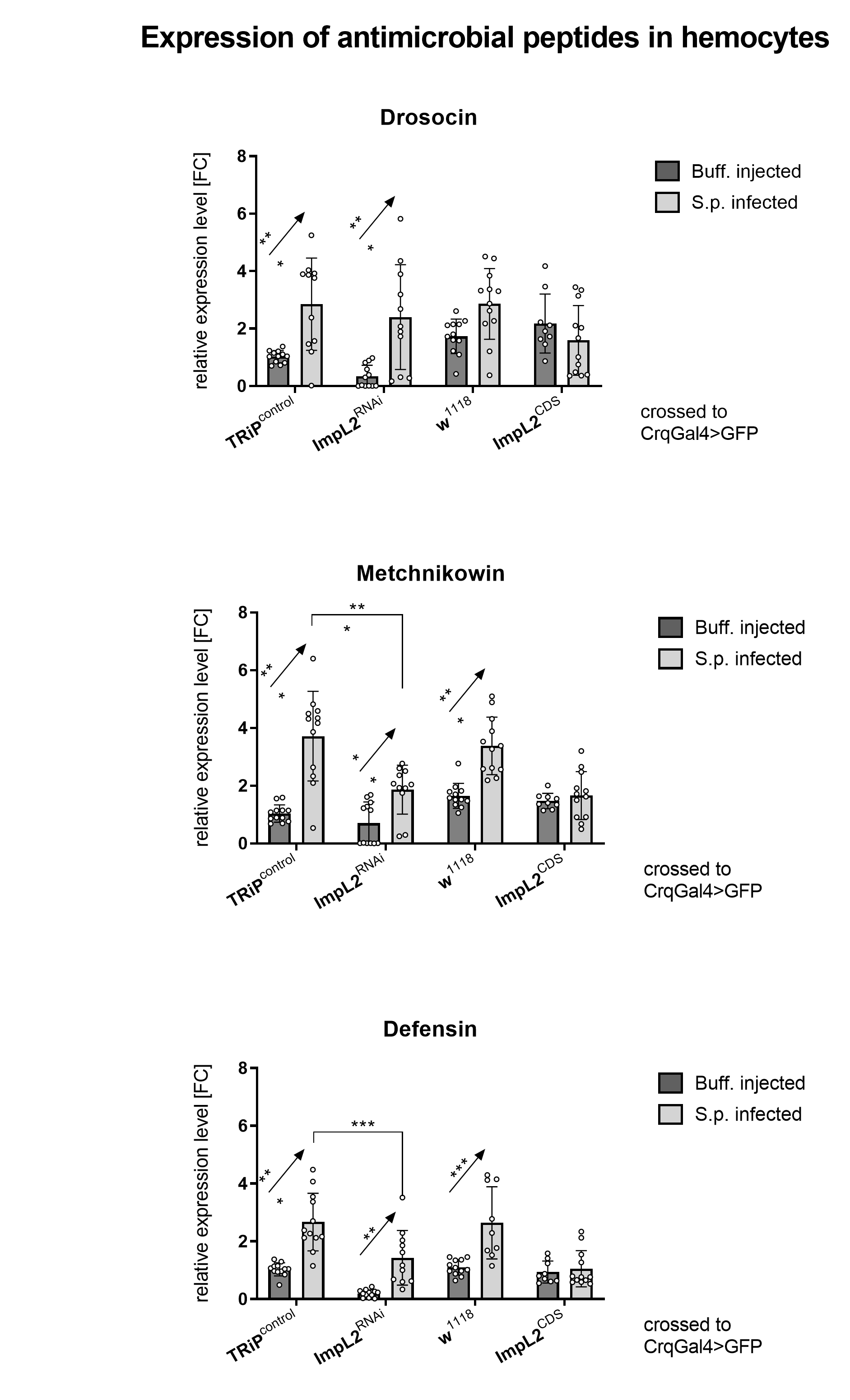
